# Mechano-signaling feedback underlies precise inner hair cell patterning in the organ of Corti

**DOI:** 10.1101/2022.01.24.477468

**Authors:** Roie Cohen, Shahar Taiber, Olga Loza, Shahar Kasirer, Shiran Woland, David Sprinzak

**Author notes:** Equally contributing authors (Roie Cohen, Shahar Taiber, Olga Loza).

## Abstract

The mammalian hearing organ, the organ of Corti, is one of the most organized tissues in the mammalian body plan. It contains precisely positioned array of alternating sensory hair cells (HC) and non-sensory supporting cells that emerge during embryonic development from an initially disordered pro-sensory domain. While much is known on the genetics and biochemistry underlying this process, it is still unclear how such precise alternating patterns emerge during embryonic development. Here, we combine live imaging of mouse inner ear explants with hybrid mechano-regulatory models to elucidate the mechanisms underlying the formation of a single row of inner HC (IHC). We show that a narrow strip of initially disordered salt-and-pepper pattern, generated by Notch-mediated lateral inhibition, is dynamically refined by coordinated intercalations, delaminations, and differential adhesion. We identify a new morphological transition, termed ‘hopping intercalation’, that allows nascent IHC to ‘hop’ under the apical plane into their final position. We further show that IHC patterning is associated with boundary localization of the cell adhesion molecules, Nectin-3 and Nectin-1. Our experimental results and modeling support a mechanism for precise patterning based on a feedback between Notch-mediated differentiation and mechanically driven cellular reorganization that is likely relevant for many developmental processes.

## Introduction

The formation of complex, yet precise, cellular patterns during embryonic development relies on two key aspects: the ability of cells to coordinate their differentiation with their neighbors through biochemical signals, and the control of cellular and tissue morphology through cell mechanics. It is often unclear, however, how these two aspects of development are coordinated to produce precise cellular patterns. An emerging concept in morphogenesis in recent years is that feedback between biochemical signals and mechanical forces, termed mechano-signaling feedback, is crucial for many developmental processes (Lenne et al., 2021; Maroudas-Sacks and Keren, 2021). However, relatively little is known on how such mechano-signaling feedback controls a large class of developmental patterning processes involving regular alternating cellular patterns, where neighboring cells acquire distinct regulatory and mechanical properties. In most research to date, formation of such alternating patterns was studied in the context of models such as Turing-type models and signaling mediated lateral inhibition models (Collier et al., 1996; Turing, 1990). Mounting evidence in multiple systems strongly suggests that in many of these examples mechanical forces also play a crucial role in coordinating developmental patterning (Cohen and Sprinzak, 2021; Mammoto and Ingber, 2010).

A prime example of an alternating cellular pattern controlled by both signaling mediated differentiation and mechanically driven reorganization is the mammalian hearing organ, the organ of Corti (OoC), located within the cochlea (Fig. 1a). The OoC, arguably the most organized tissue in the mammalian body plan, consists of exactly four rows of sensory hair cells (HC), interspersed by non-sensory supporting cells (SC), to create an alternating checkerboard-like pattern (Fig. 1a). These four rows are further divided into one row of inner hair cells (IHC) and three rows of outer hair cells (OHC), with a single line of inner pillar cells (PC) separating them. This precise organization of sensory HC is believed to have a crucial role in hearing sensitivity and frequency selectivity in mammals (Huang et al., 2012; Manoussaki et al., 2006).

**Figure 1.**
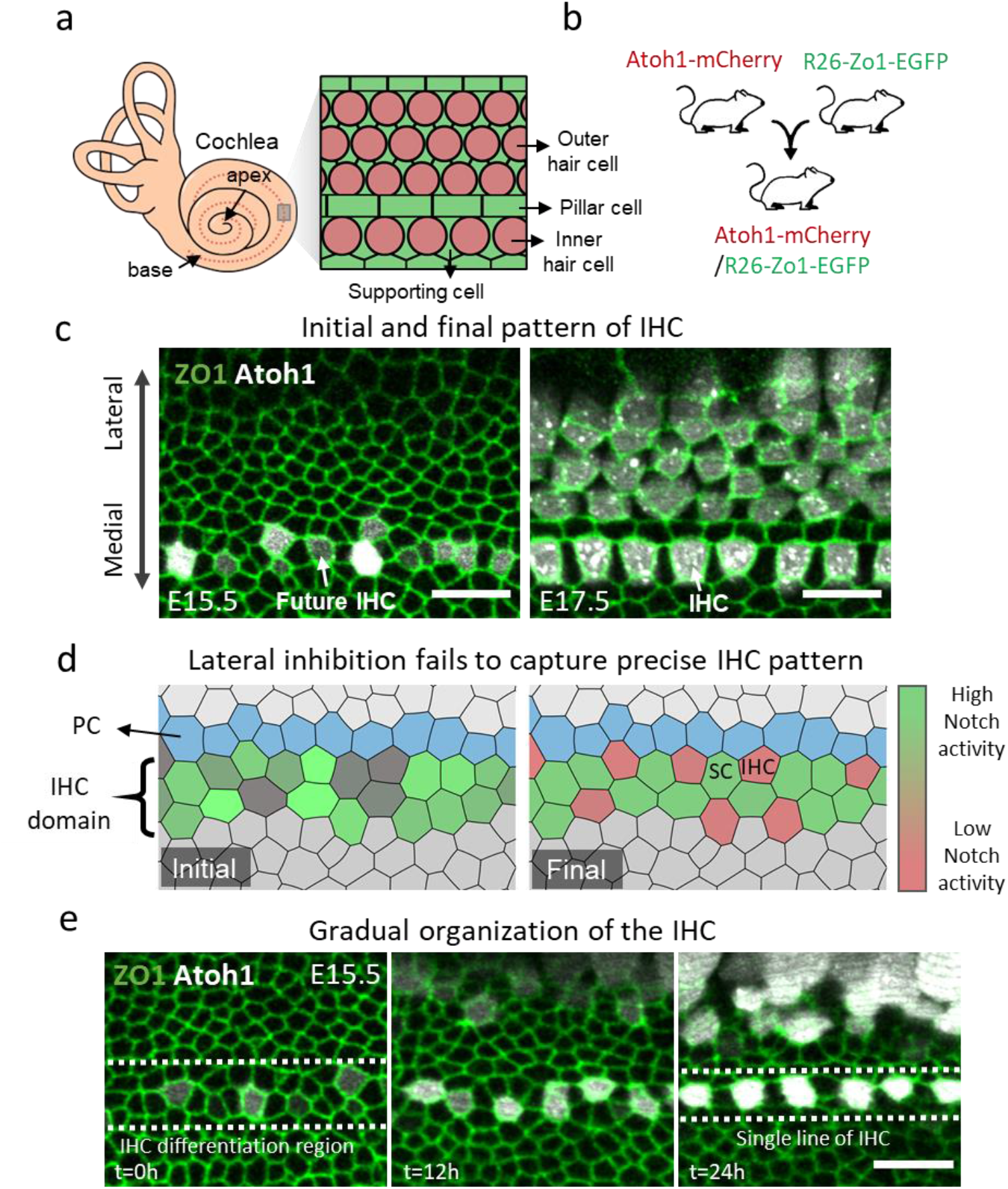
A single row of IHC is formed from an initially disordered salt-and-pepper pattern. **(a)** Schematic of the mammalian inner ear. Dashed line represents the location of the OoC along the cochlea. A zoom-in on the OoC with marked OHC, IHC, SC and PC is presented on the right. **(b)** Schematic of the transgenic mice used for imaging. Atoh1-mCherry (HC marker) mice are crossed with ZO1-EGFP (tight junction marker) mice to produce double-reporter mice. **(c)** Snapshots from movies of cochlear explant of a ZO1-EGFP (green)/Atoh1-mCherry (gray) mouse showing initial (left, E15.5) and advanced (right, E17.5) patterns of the IHC row. Double arrow displays the medial-lateral axis. **(d)** Snapshots from the initial (left) and final (right) timepoints of a lateral inhibition simulation over a narrow strip of cells (Movie S2). Color bar represents Notch activity level. Initial pattern is resolved to a salt-and-pepper pattern of low Notch activity HC (red) and high Notch activity SC (green). **(e)** A filmstrip from a movie of an E15.5 cochlear explant showing the gradual differentiation and patterning of the IHC row (Movie S3). Scale bar in (c) and (e): 10μm.

The OoC initially develops from a disordered undifferentiated prosensory epithelium and gradually differentiates and organizes into an ordered pattern (Driver and Kelley, 2020). In mice, HC differentiation begins after cells in the OoC exit cell cycle at around embryonic day 14 (E14) and reaches a fully organized pattern by postnatal day 0 (P0). HC differentiation is associated with the expression of the transcription factor Atoh1, which is considered to be the master regulator of HC fate and is also one of the earliest markers of differentiation into HC (Brown and Groves, 2020). During its development, the OoC also exhibits a spatial developmental gradient along the base-to-apex axis of the cochlea, where the base and the apex are the most and least developed regions, respectively.

The alternating pattern of HC and SC in the OoC emerges through a process of Notch-mediated lateral inhibition (Brown and Groves, 2020). In lateral inhibition, cells compete to differentiate into a primary cell fate (e.g. HC in the OoC) by inhibiting their neighbors from also adopting the same fate (Binshtok and Sprinzak, 2018). Thus, within a small group of cells, one cell adopts a primary fate, while all its neighbors adopt the secondary fate (e.g. SC in the OoC), leading to an alternating salt-and-pepper pattern. The inhibition is typically mediated by the Notch signaling pathway, which is a conserved juxtacrine signaling pathway driving communication between neighboring cells (Henrique and Schweisguth, 2019; Kovall et al., 2017; Siebel and Lendahl, 2017; Sprinzak and Blacklow, 2021). While lateral inhibition models can generate different types of alternating patterns in either 1D or 2D cell layers, it cannot produce by itself highly organized patterns when applied to disordered cell lattices (Cohen et al., 2020; Shaya et al., 2017). Thus, the highly organized pattern of HC and SC in the OoC cannot be solely explained by simple lateral inhibition models.

In a recent study, we used live imaging of inner ear explants and mathematical modeling to show that the precise patterning of the OHC region in the OoC emerges from an initially disordered salt-and-pepper pattern by reorganization processes driven by mechanical forces (Cohen et al., 2020). We have shown that external shear and compression forces acting on the OHC region and repulsion forces between HC can drive the transition from a disordered to ordered pattern. The pattern is further refined by differential adhesion between OHC and SC. Thus, the precise pattern of OHC is not simply a result of signaling mediated differentiation (e.g. lateral inhibition), but rather relies on mechanical forces that rearrange the cells into their precise final pattern.

In this work, we explore how the feedback between Notch-mediated lateral inhibition and mechanical forces gives rise to the formation of a single row of IHC, exhibiting a precise alternating pattern of IHC and SC. We used live imaging of inner ear explants from transgenic mice expressing both ZO1-EGFP and Atoh1-mCherry (Fig. 1b), allowing us to track the morphological dynamics of the cells and their differentiation state, respectively. We found that an initially disordered salt-and-pepper pattern spanning 2-3 cell rows is refined by coordinated intercalations and delaminations. In particular, we identified a new type of intercalation termed ‘hopping intercalation’, allowing Atoh1+ cells to ‘hop’ under the apical plane towards the PC row. We also found that some out-of-line cells expressing low levels of Atoh1 delaminate. We developed a hybrid modeling approach combining a lateral inhibition model with a mechanical 2D vertex model that captures the main features observed experimentally. We show that further morphological features can be explained by differential adhesion between IHC, SC, and PC. Predictions generated by our model are validated by tracking rearrangements following laser ablation of single cells. Finally, we found that localization of Nectin-3 and Nectin-1 at the IHC:PC boundary correlates with the onset of IHC patterning process suggesting a role for Nectins in refinement of IHC patterning. Overall, these results support a picture where mechano-signaling feedback coordinates the precise alternating pattern of IHC.

## Results

### Inner hair cells differentiate and reorganize into a single row between E15.5 and E17.5

The IHC row in the postnatal cochlea exhibits a precise long-range alternating pattern with very few defects, typically in the form of extra IHC (Fig. S1a). To examine how such a precise pattern emerges during development, we used an assay based on live imaging of cochlear explants allowing tracking of both cell morphology and differentiation state of cells. This was performed by timed mating of transgenic mice expressing ZO1-EGFP, marking the tight junctions at the apical surface of the tissue, with transgenic mice that express a transcriptional reporter, Atoh1-mCherry, which is one of the earliest markers for differentiation into nascent HC (Chen et al., 2002; Lepelletier et al., 2013)(Fig. 1b, Fig. S1b). Cochleae from these double transgenic mice were dissected at different developmental stages (E15.5 and E17.5) and imaged under an incubated microscope for up to 24 hours.

Time lapse movies of cochlear explants at E15.5 allows tracking of the initial stages of differentiation as observed by the emergence of Atoh1+ cells (Movie S1). Consistent with previous work (Driver and Kelley, 2020), this differentiation begins at the base and progresses towards the apex, with IHC emerging prior to OHC. Here, we focused on the differentiation of the IHC row. Initial appearance of Atoh1+ cells at the apex region of E15.5 cochleae occurs within a 2-3 cell rows wide strip at the medial prosensory domain (Fig. 1c, left). By E17.5, Atoh1+ positive cells are organized as one ordered row throughout most of the cochlea (Fig. 1c, right), similar to the pattern observed in the adult ear.

The differentiation into HC and SC fates is driven by the process of Notch-mediated lateral inhibition (Brown and Groves, 2020). To test whether lateral inhibition can create an ordered pattern of IHC from a narrow strip of cells, we applied our previously described contact-based model of lateral inhibition (Formosa-Jordan and Sprinzak, 2014; Shaya et al., 2017). For simplicity, we consider only one type of Notch ligand in the model, although both Dll1 and Jag2 have been shown to participate in this process (Kiernan et al., 2005). In these simulations we also assumed boundary conditions where the cells in the medial region to the differentiating cells (Kolliker’s organ) express low levels of Notch ligands. This is based on the observation that cells in the Kolliker’s organ express Jag1 (Basch et al., 2016; Orvis et al., 2021). Running our model on a narrow strip of cells results in a salt-and-pepper pattern of HC surrounded by SC, albeit without a particular long-range order (Fig. 1d, Movie S2). The salt-and-pepper patterns we obtained in our basic lateral inhibition models suggest that lateral inhibition is not sufficient to explain the highly organized pattern of the IHC. As we have previously shown for the OHC region (Cohen et al., 2020), cellular reorganization processes may contribute to formation of a precise pattern. We therefore wanted to focus on the dynamics of IHC differentiation using live imaging of inner ear explants at this developmental stage.

To observe the reorganization processes that occur during the patterning of the IHC row, we obtained movies of inner ear explants at early developmental stages. Live imaging of E15.5 cochleae at the apex region, showed that an initially disordered salt-and-pepper pattern of Atoh1+ cells over 2-3 cell rows reorganize into an aligned single row of Atoh1+ cells (next to the future PC row) within ~1 day (Fig. 1e, Movie S3). Analysis of these movies revealed local morphological transitions and reorganization processes that contribute to the formation of a single row. We found that Atoh1+ cells that do not touch the future PC row move laterally towards the PC row soon after their differentiation (Fig. S1c). While we cannot determine what drives this lateral motion, we observed frequent cell divisions in the Kölliker’s organ, medial to the developing IHC domain, in agreement with a previous study showing increased cell divisions in the medial part of the cochlear duct (Fig. S1d, Movie S4) (Ishii et al., 2021). This might suggest that a pressure gradient is generated in the medial-lateral axis, compressing the Atoh1+ cells towards the lateral prosensory domain. Such a pressure gradient is expected to act on all the cells and not only on Atoh1+ cells, however, it was previously shown that cells that upregulate Atoh1 detach from the basement membrane and migrate towards the luminal surface (Tateya et al., 2019). This could explain the high mobility of Atoh1+ cells relative to other cells that are still connected to the basement membrane.

### Atoh1+ cells perform ‘hopping intercalations’ towards the PC border

Interestingly, we found that the lateral movement of nascent Atoh1+ cells is accompanied by two morphological transitions: (1) cellular intercalations as previously shown to occur during the development of the OoC (Cohen et al., 2020; Driver et al., 2017)(Fig. S2a-d, movie S5), and (2), a newly identified reorganization process, termed ‘hopping intercalation’. Hopping intercalation occurs when a cell opens a new apical footprint at an adjacent tri-cellular junction and then merges with the original apical surface (Fig. 2a-d, movie S6). In this process, the cell effectively ‘hops over’ the cellular junction instead of squeezing in as occurs in standard intercalation. To verify that hopping intercalation has a significant role in the patterning process, we have counted the number of intercalations in our movies and found that hopping intercalations account for roughly 40% of the total intercalations towards the future PC border. Given that a cell needs to deform its apical surface in order to squeeze in between two cells, an Atoh1+ cell might initiate hopping intercalation to reduce its mechanical energy. This is consistent with our previous observation that the apical surface of HC is less deformable than that of SC (Cohen et al., 2020). A supporting indication for this hypothesis is the observation that the apical surface area of Atoh1+ cells after hopping intercalation is significantly larger than the surface area before the hopping intercalation (Fig. 2e).

**Figure 2.**
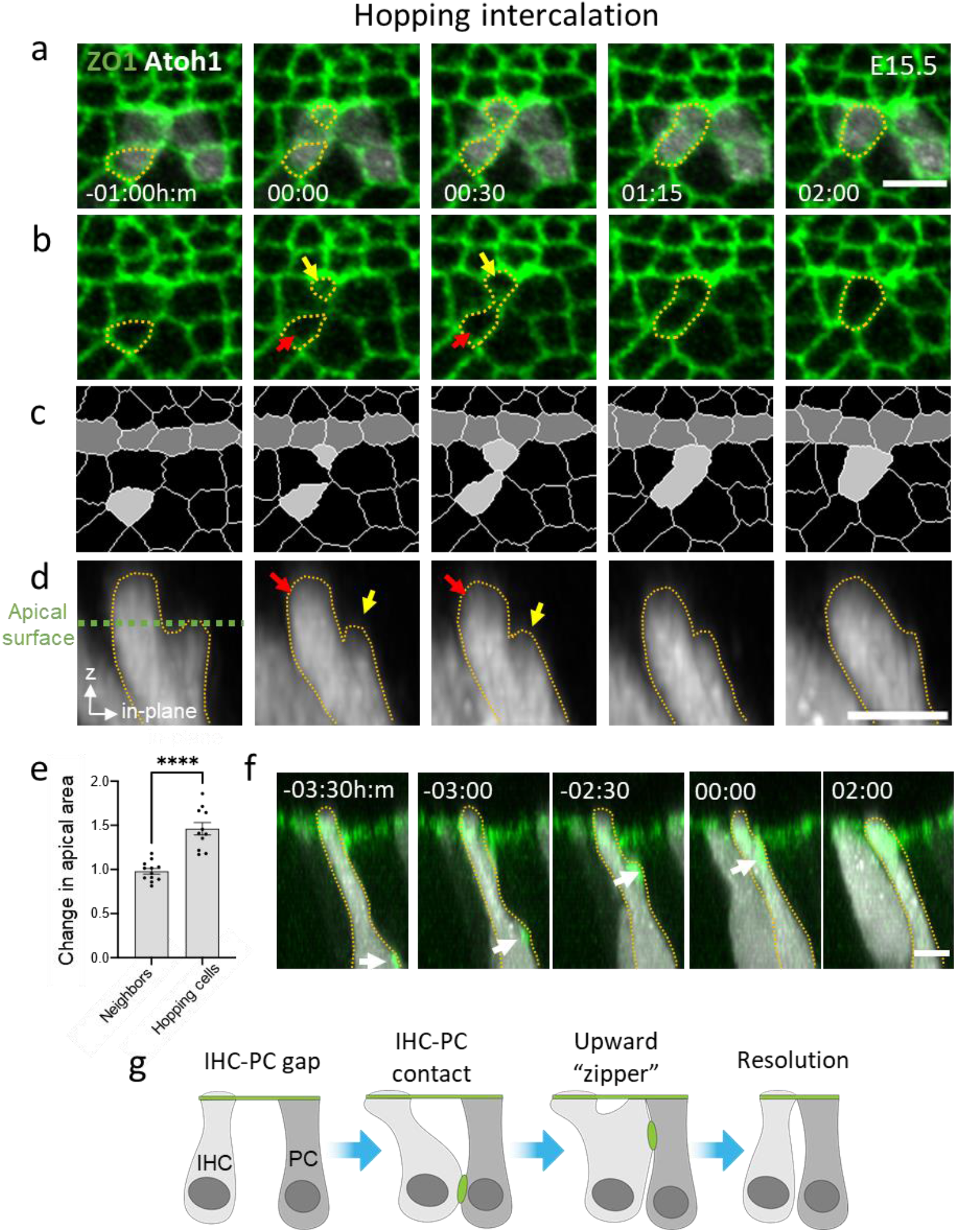
Hopping intercalation of nascent IHC towards the PC row. **(a-c)** A filmstrip from an E15.5 cochlear explant showing an apical view of a hopping intercalation event with (a) both Atoh1 and ZO1 markers (b) only the ZO1 marker, and (c) a segmentation of (b) highlighting intercalating cell (light gray) and PC (dark gray) (Movie S6). New apical surface opens up (yellow arrow in (b)) and merges with the original apical surface (red arrow in (b)). **(d)** A cross section showing a side view of the intercalating cell. Dashed line marks the apical surface of the tissue. Red and yellow arrows mark the original and new apical surfaces of the cell, respectively. **(e)** Change in apical area of hopping cells. Graph shows the relative change in apical area (area after/area before hopping) for hopping cells. For control, we measured the change in apical area of non-hopping neighboring IHC at similar times. **(f)** A cross section of the filmstrip in (a-c) (zoomed out with respect to (d)) showing a ZO1 punctum propagating towards the apical surface. Timestamps show time with respect to the initiation of the hopping event (at 00:00h:m). **(g)** A cross section schematic of a hopping intercalation. Statistical test in (e): Welch’s t-test, plot shows mean ± SEM. n=12, 11 repeats of neighbors and hopping cells, respectively. ****P<0.0001. Scale bars in (a), (d) and (f): 5μm

To better understand the mechanics of hopping intercalation, we examined the 3D morphology of the Atoh1+ cells during and before hopping. We found that several hours prior to hopping, the cell body below the apical surface shifts laterally towards the PC row and seems to be in contact with it before a new apical surface emerges (Fig. 2f). To make sure this shift is not an artifact of the ex-vivo culture, we additionally examined fixed samples of E15.5 cochleae and observed a similar lateral shift of Atoh1+ cells in the apex region (Fig. S2e). While the cause for the lateral shift is unknown, this could be a result of the lateral compression discussed previously. Additionally, in some cases of hopping intercalations (but also in some standard intercalations), we observed a sub-apical ZO1 punctum at the sub-apical contact point between the Atoh1+ cell and the future PC. This punctum migrates towards the apical surface prior to the initiation of intercalation and seems to attach the Atoh1+ cell to the future PC in a zipper-like motion (Fig. 2f, g, Fig. S2d). As far as we know, this is the first observation of hopping intercalation. This type of intercalation expands the set of morphological transitions capable of reorganizing epithelial layers.

### A hybrid regulatory-mechanical model of hopping intercalations captures IHC patterning

The observation that nascent IHC are pushed laterally, resulting in intercalations and hopping intercalations towards the PC border may provide a potential explanation for how Atoh1+ cells align into a single row. To understand whether these observations are sufficient to explain the patterning of the IHC row, we developed a hybrid model that incorporates cellular dynamics with a lateral inhibition model (Fig. 3a). To model cellular dynamics, we used a modified 2D vertex model, a well-established approach for simulating dynamics in epithelial tissues (Farhadifar et al., 2007). In a 2D vertex model each cell and each cellular boundary can be assigned with different mechanical properties, and the total mechanical energy for the tissue can be defined (Fig. S3a). Given an initial state with a predefined set of mechanical parameters, the model modifies the configuration of the vertices such that the total mechanical energy is minimized. In a basic 2D vertex model, each tri-cellular junction is represented by a vertex and each cell is represented by a polygon with edges that connect the vertices associated with that cell. To also simulate the curvature of the cellular boundaries, we extended the basic 2D vertex model by introducing additional degrees of freedom in the form of virtual vertices on each edge (Fig. S3b and methods) (Frost et al., 1988; Tamaki et al., 2012). The modified 2D vertex can account for non-polygonal cell shapes (e.g. rounder HC), as well as morphological transitions involving non-polygonal cellular deformations.

**Figure 3.**
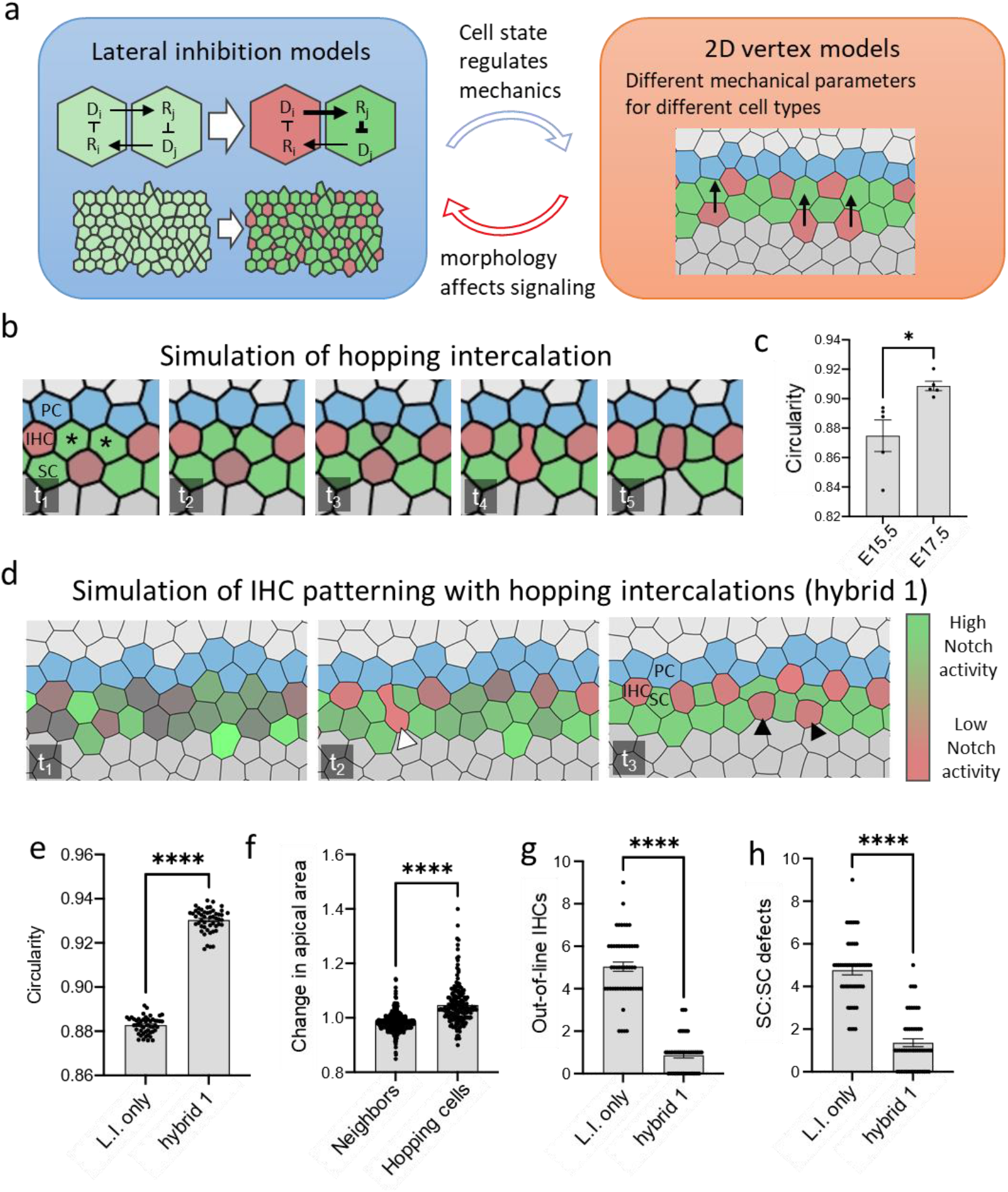
A hybrid modeling approach incorporating hopping intercalations accounts for reduction in patterning defects. **(a)** Schematic of the hybrid modeling approach incorporating lateral inhibition, cell mechanics, and the feedback between them. A simplified circuit of lateral inhibition feedback and the resulting pattern is shown on the left panel (see methods for full description). 2D vertex models (right panel) are used to simulate how mechanical forces (arrows) drive morphological transitions. **(b)** A filmstrip of a simulation showing hopping intercalation of a HC towards the PC row. Asterisks mark a SC:SC defect being resolved by hopping intercalation (see methods for detailed description) (Movie S7). **(c)** Comparison between circularity of IHC at E15.5 and E17.5. Circularity is defined as 4*πA/L*^2^, where A and L are the apical area and circumference of the cell, respectively. **(d)** A filmstrip of a simulation of the hybrid model combining lateral inhibition and hopping intercalations (see hybrid1 model in Table 1) (Movie S8). White arrowhead shows an intercalating cell. Black arrowheads show out-of-line IHC defects. **(e-h)** Quantitative comparison between a model with lateral inhibition only (L.I. only) and a hybrid model of lateral inhibition and hopping intercalations (hybrid 1 model, see methods for details). Circularity of IHC (e), number of out-of-line IHC defects (g), and number of SC:SC defects (h) are shown for the two models. Panel (f) shows relative change in apical area (area after/area before hopping) for hopping cells in hybrid 1 model. For control, we measured the change in apical area of non-hopping neighboring IHC in the simulation. Statistics: (c), (e) and (f), Welch’s t-test, (g) and (h), Mann-Whitney test, all plots show mean ± SEM. Repeats: (c) n=5 independent movies for each time point, (e) n=50, (f) n=175, 457 for hopping cells and neighbors, respectively, (g-h) n=50 simulations. ****P<0.0001, *P<0.05.

As hopping intercalation is a newly observed morphological transition, we wanted to find a way to model it such that we capture the qualitative behavior observed experimentally. Hopping intercalation begins with the opening of the second apical surface of the hopping cell and ends at the merger of the two apical surfaces. We model this process as follows: (1) The second apical surface is modeled as an individual cell that is linked to the original cell through the area energy term (Fig. S3a). (2) As the opening of a new apical surface requires an active force to push on the neighboring cells (potentially originated by the cell body “pushing up”), an additional expansion energy term is assigned for the cell representing the new apical surface. (3) Once the two apical surfaces get close and intercalate, they immediately merge into a single cell. Modeling hopping intercalation in such a way creates a similar hopping process to the one observed experimentally (Fig. 3b, Movie S7).

To simulate the effect of hopping intercalations on IHC patterning, we applied a hybrid model combining lateral inhibition and a 2D vertex model that includes hopping intercalation. The two parts of the model are advanced in time in a synchronous manner, allowing for differentiation and morphological dynamics to progress in parallel (see methods). Based on our observations, the assumptions that we make in the 2D vertex model are the following: (1) Atoh1+ cells have higher contractility and incompressibility. This assumption is based on the observed higher circularity of HC relative to SC (Fig. 3c) and consistent with our previous observation that HC are less deformable than SC (Cohen et al., 2020). (2) We assume local repulsion force between HC. This is based on our previous observations that HC physically touch each other at the sub-apical plane corresponding to the position of the nuclei (Cohen et al., 2020). (3) Atoh1+ cells are subjected to a lateral compression force towards the PC. This force is applied on all cells whose Atoh1 level crosses a certain threshold. This assumption is based on the observed lateral motion of HC towards the PC row (Fig. S1c). (4) Hopping intercalation is initiated for Atoh1+ cells whose area decreases below a certain threshold due to the lateral compression (see methods). Applying these assumptions to our model (termed hybrid 1 model) captures the lateral movement of Atoh1+ cells and hopping intercalations as observed experimentally (Fig. 3d, Movie S8). In terms of HC morphology, the hybrid 1 model captures the increase in circularity of HC (Fig. 3e), as well as the increase in area following the hopping intercalation events (Fig. 3f).

To understand whether the hybrid 1 model generates a more ordered pattern of IHC, we examined the defects in the resulting pattern. We identified two types of defects that occur during the patterning process: The formation of extra HC that do not border the PC row (termed out-of-line IHC defects, black arrowheads in Fig. 3d), and the presence of two neighboring SC with no HC between them (termed SC:SC defects, asterisks in Fig. 3b). We have compared the number of defects between a model of lateral inhibition without hopping intercalation (L.I. only model) and the hybrid 1 model that contains hopping intercalation. We find a significant reduction in both types of defects in the hybrid 1 model (Fig. 3g, h). Thus, we conclude that hopping intercalations helps reduce the number of defects and drive the pattern of IHC to a more organized state. We note, however, that even with hopping intercalations there are still defects of both types that are not resolved in our model, suggesting the presence of additional mechanisms for defect resolution.

### Cells with low Atoh1 levels delaminate

In addition to the intercalations and hopping intercalations, we also noticed that some of the Atoh1+ cells that are not aligned against the future PC delaminate (Fig. 4a, Movie S9). Such delaminations contribute to reducing the number of out-of-line IHC. To determine what causes these delaminations, we tested if there is a correlation between the decision of a cell to delaminate, its Atoh1 level, and its position relative to the PC row. We find that delaminating Atoh1+ cells have, on average, a significantly lower expression level of Atoh1 relative to their non-delaminating Atoh1+ neighbors (Fig. 4b). Furthermore, most delaminating Atoh1+ cells are positioned more medially to non-delaminating Atoh1+ cells, and do not touch the future PC row (Fig. 3c). Correspondingly, we find that cells that do not touch the PC row typically have lower Atoh1 expression (Fig. 3d). We do occasionally observe weak Atoh1+ cells that do not touch the PC row and do not delaminate. It is reasonable to assume, however, that these defects are either resolved through delaminations at a later time point not captured in our movies, or do not resolve and persist as the sporadic defects observed in the final pattern (Fig S1a). These observations provide a potential mechanism for resolving out-of-line IHC defects and relates the decision to delaminate with a reduced level of Atoh1. We note that we were not able to track and resolve the fate of the delaminating cells in our current setup. It has been shown that cells in the sensory epithelium forced to inactivate the expression of Atoh1 undergo apoptosis, suggesting that an apoptotic program is associated with cells ceasing to express Atoh1 (Cai et al., 2013).

**Figure 4.**
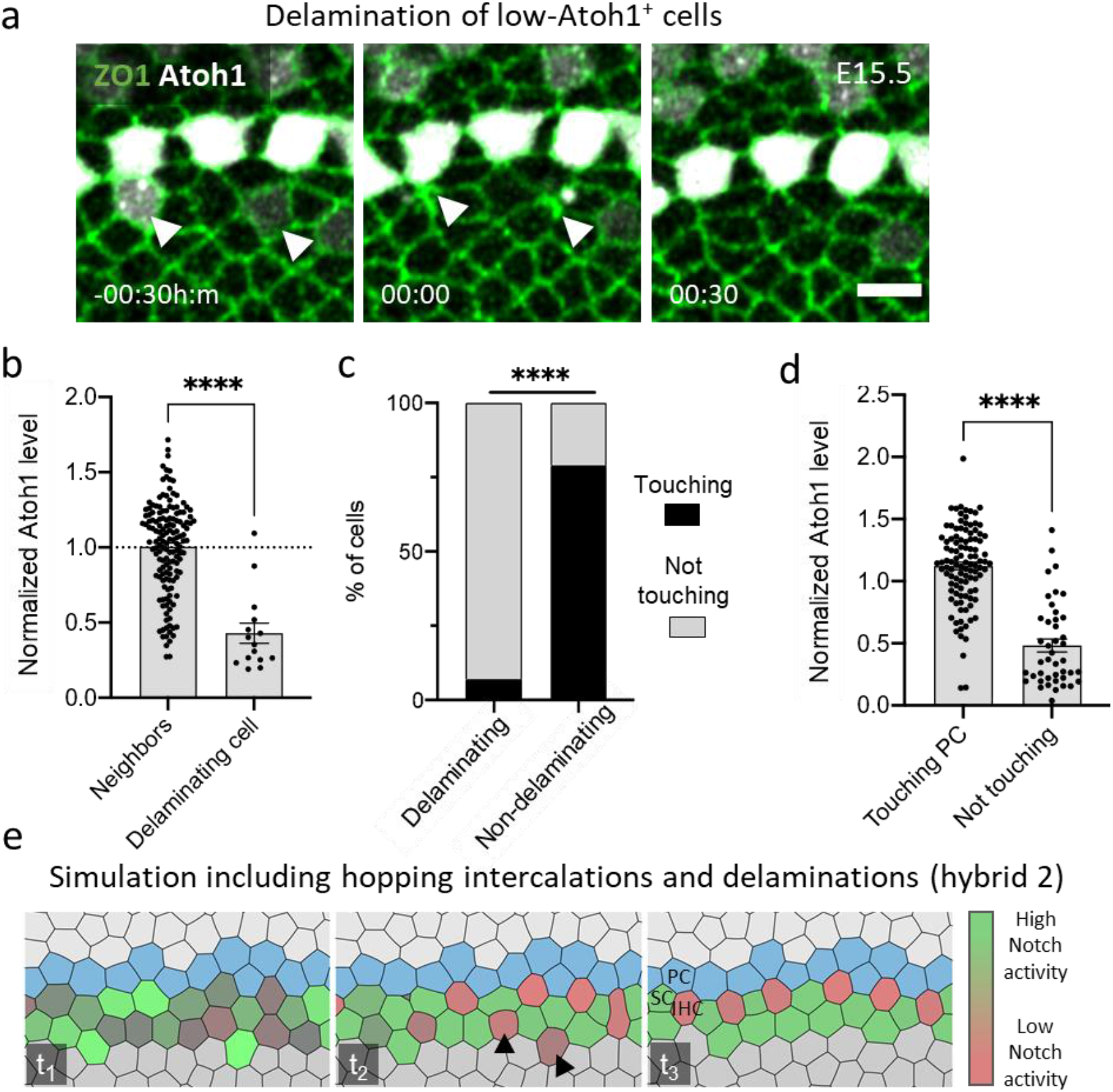
The IHC pattern is refined by delaminations of low Atoh1+ cells. **(a)** A filmstrip showing delaminations (arrowheads) of low-Atoh1^+^ cells (Movie S9). **(b)** Comparison of normalized Atoh1 levels in delaminating Atoh1+ cells vs. non-delaminating Atoh1+ cells. **(c)** Comparison of the percentage of IHC that touch\do not touch the PC row for delaminating and non-delaminating Atoh1+ cells. **(d)** Normalized Atoh1 level in HC that touch the PC row relative to HC that do not touch the PC row. **(e)** Simulation of the patterning process of IHC that incorporates both hopping intercalation and delaminations of IHC that do not touch the PC row (Hybrid 2 model) (Movie S10). Arrowheads mark two out-of-line IHC defects that were resolved by the end of the simulation. Statistics: (b) and (d), Mann-Whitney test, plots show mean ± SEM. (c) Chi-square test. Repeats: (b) n= 15 delaminating cells and 150 neighbors, (c) n=15, (d) n=43, 106 for cells not touching the PC row and cells touching, respectively. ****P<0.0001. Scale bar in (a): 5μm

Based on these observations, we added a simplified delamination rule to the hybrid 1 model such that each HC that does not manage to border the PC row within a certain time frame is forced to delaminate (termed hybrid 2 model). Not surprisingly, adding this delamination rule to the model completely removes the out of line HC defects and produces a more organized alternating pattern of IHC (Fig. 4e, Movie S10). While more complex assumptions associating Atoh1 activity with delaminations may provide a more realistic model, we do not currently have sufficient information (e.g spatial dependence of Notch activity) to incorporate such associations in our model.

### Notch inhibition leads to extra Atoh1+ cells, but delaminations persist

It has been previously shown that inhibition of Notch signaling leads to the formation of extra HC (Mizutari et al., 2013; Yamamoto et al., 2006). While complete Notch inhibition is known to lead to massive production of HC and complete disarray of the HC pattern, partial Notch inhibition leads to more subtle phenotypes exhibiting milder addition of IHC (Basch et al., 2016). We therefore wanted to test how the dynamics of organization are affected by such partial Notch inhibition. We performed live imaging of explants in the presence of sub-saturating levels of DAPT, a γ-secretase inhibitor, (Fig S4a, Movie S11) and tracked the observed dynamics. We found that partial Notch inhibition leads to generation of additional Atoh1+ cells within 12-24 hours after adding DAPT (Fig S4b-c). We found however, that some weak Atoh1+ cells still delaminate even under such inhibition (Fig S4d). Hence, partial inhibition of Notch signaling leads to generation of extra Atoh1+ cells but is not sufficient to prevent delaminations.

### Differential adhesion refines IHC pattern

The model that includes both intercalations and delaminations (hybrid 2 model) can explain the precise alternating order of IHC and SC. However, it fails to capture additional morphological features including the increase in alignment of PC, and the longer contact length between IHC and PC (IHC:PC contact) compared to the contact length between SC and PC (SC:PC contact) (Fig. 5a-c). We reasoned that these morphological aspects may be due to differences in adhesion between the different cell types that dynamically emerge during development. To introduce these changes in the model, we added additional assumptions on the line tensions associated with the IHC:PC, IHC:SC and IHC:IHC boundaries (termed hybrid 3 model) (Fig. S3a). For simplification, these changes were introduced as a second stage in the simulation, immediately after reaching the final pattern in the hybrid 2 model (Note that these two stages likely overlap *in vivo*). The adhesion rules we introduced are: (i) Tensions between PC and other cell types (e.g. IHC, SC) are higher than tensions between PC and other PC. Such high tension on the boundary between domains has been shown to promote more straight boundaries in other systems (Aliee et al., 2012). (ii) We assumed that the tension on IHC:PC boundary is lower than the tension on SC:PC boundary. This assumption is based on the observation that the IHC:PC contact length is longer than that of the SC:PC contact length (Fig. 5c). (iii) We assumed very high tension between IHC. This is based on the observation that in the infrequent occasion that two Atoh1+ cells are in contact, the length of the contacts are typically very small (see arrowheads in Fig. S4a). Simulations of the hybrid 3 model that includes adhesion rules captured the increase in alignment of PC (Fig. 5d,e) and the longer IHC:PC contact length compared to SC:PC boundary (Fig. 5d,f) (Movie S12). We note that longer IHC:PC contacts are already observed at early stages (E15.5) suggesting that the changes in the tension of the IHC:PC occur prior to changes in tensions on other boundaries. Overall, the results from hybrid 3 model show that addition of adhesion rules between different cell types helps refine the pattern of the IHC row.

**Figure 5.**
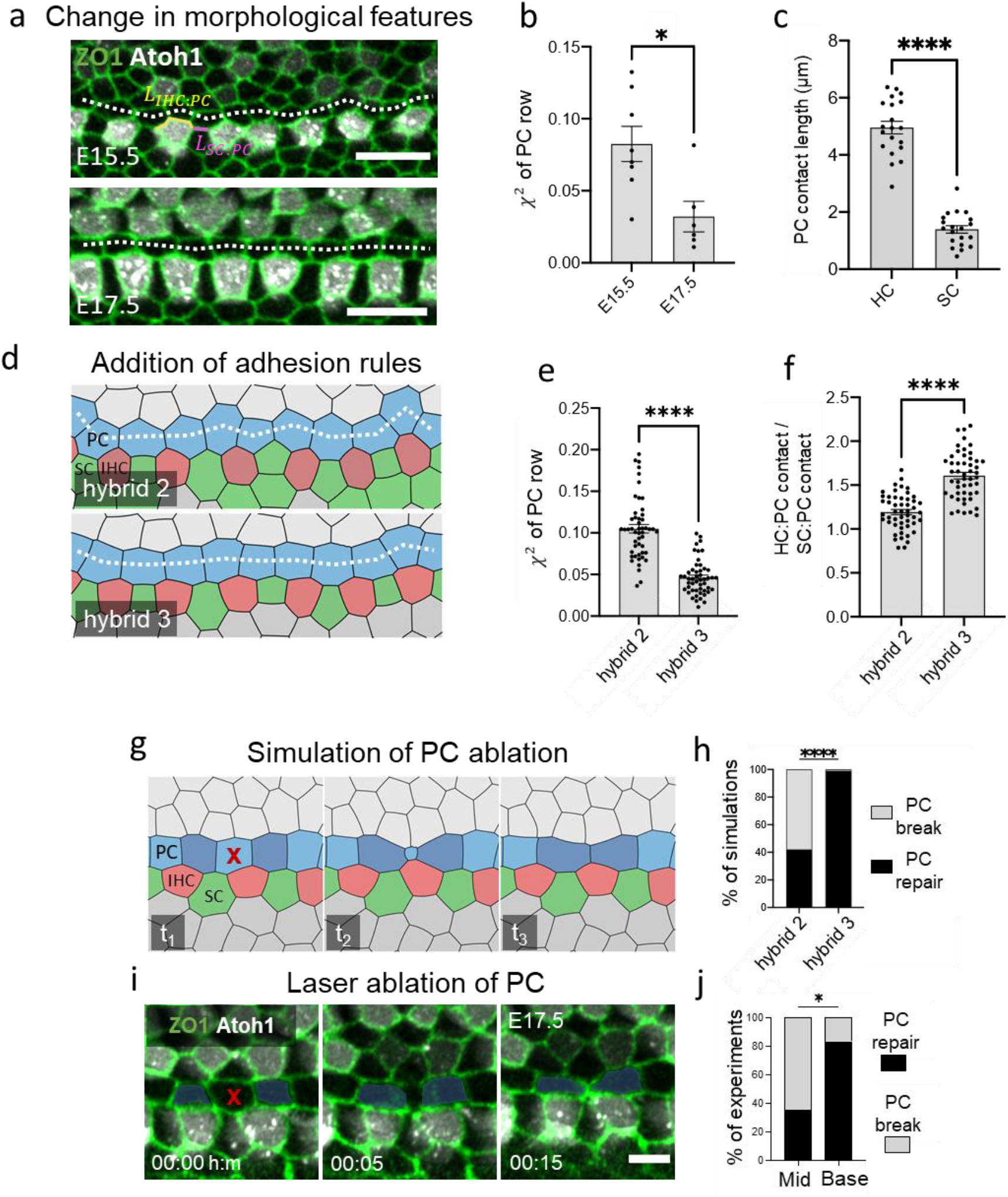
Differential adhesion underlies straightening of IHC:PC boundary. **(a)** Two images from E15.5 and E17.5 cochlear explant movies showing the increase in straightness of the IHC and PC rows (dashed line connects the centers of PC). **(b)** Quantification of alignment of PC row at E15.5 and E17.5 measured by the reduced *χ^2^* (see methods). **(c)** Comparison of the mean length of HC:PC and SC:PC contact at E15.5. **(d)** Comparison between hybrid models that include (hybrid 3) or do not include (hybrid 2) adhesion rules. For simplification, adhesion rules in hybrid 3 model are added after final pattern is generated by the hybrid 2 model (Movie S12). Hybrid 3 model includes differences in line tensions such that tension(PC:SC)=tension(HC:HC)>tension(PC:HC)>tension(PC:PC) (see Fig. S3 and methods). **(e)** Quantification of alignment of PC row for the two models measured by the reduced *χ*^2^. **(f)** Ratio between the mean length of HC:PC and SC:PC boundaries in the two models. **(g)** Filmstrip of a simulation from the hybrid 3 model, where a single PC was ablated (red ‘x’) (Movie S13). The gap created by the ablated PC is repaired by the neighboring PC. **(h)** Fraction of PC ablation simulations that resulted in the repair of the PC row by neighboring PC in the two models. **(i)** A filmstrip from a laser ablation experiment performed at the base region of an E17.5 cochlea, where a single PC was ablated (red ‘x’) (Movie S15). Similar to the simulation in (f), the gap was repaired by the neighboring PC. **(j)** Fraction of PC ablation experiments that resulted in the repair of the PC row by neighboring PC. Statistics: (b) and (f), student’s t-test, (c) Welch’s t-test, (e) Mann-Whitney test, plots show mean ± SEM. (h) and (j), Chi-square test. Repeats: (b) n=7, 6 for E15.5 and E17.5, respectively, (c) 20 boundaries of each type from 4 independent movies (e-f) n=50, (h) n=150, (j) n=17, 12 in the mid and base, respectively. ****P<0.0001, *P<0.05. Scale bars: (a) 10μm, (i) 5μm.

### Laser ablation experiments support differential adhesion model

To test the validity of the adhesion rules in the hybrid 3 model, we wanted to generate predictions that can be tested experimentally by applying mechanical perturbations. Local mechanical perturbations can be introduced by ablating single cells and tracking the morphological response around the ablated cell (Cohen et al., 2020). We found that simulating ablation of PC, with and without adhesion rules, results in different outcomes (see methods). In the hybrid 3 model (with adhesion rules), the PC row is predominantly repaired by the two PC neighboring the ablated cell (Fig. 5g, Movie S13). This occurs because a delaminating PC is more strongly attached to its PC neighbors, and hence ‘pulls’ on them as it delaminates. In contrast, in the hybrid 2 model (without adhesion rules), the probability of repair by the neighboring PC is about the same as the probability of a break forming in the PC row (neighboring SC moving in instead) (Fig. 5h, Fig S5a, Movie S14). To test this prediction, we performed UV laser ablation of a single PC in E17.5 cochlear explants at different positions along the base-to-apex axis, corresponding to different developmental stages. We find that ablation of PC in the base region (more advanced stage) mostly results in the repair of the PC row by neighboring PC (Fig. 5i, Movie S15). In contrast, ablation in the mid region (less advanced stage) results in a significantly smaller rate of repairs (Fig. 5j, Fig. S5b, Movie S16). These results support our hypothesis that the adhesion between PC is stronger than the adhesion to their neighbors.

### Nectins accumulate at the boundaries between IHC and PC

The adhesion rules described above, suggest the presence of dedicated molecular mechanisms that promote differential adhesion between different cell types. Earlier work has shown that differential adhesion in the OoC could be mediated by cell adhesion molecules from the Nectin family, known to promote heterophilic interactions. It was shown that Nectin-1 and Nectin-3 are differentially expressed in HC and PC, respectively, and that knockout of Nectin-1 or Nectin-3 result in patterning defects. To determine the distribution of Nectins in the relevant developmental window where IHC patterning is established, we immunostained E15.5 cochleae for Nectin-1 and Nectin-3. We observed the emergence of a polarized accumulation of Nectin-1 and Nectin-3 at the boundary between IHC and PC (Fig. 6, Fig. S6). Nectin accumulation correlates with the emergence of the IHC pattern, showing stronger accumulation at the IHC:PC boundary at the more developmentally advanced base region. This observation may suggest that the preferred adhesion of IHC to PC is mediated by heterophilic Nectin interactions. Interestingly, we also identified several puncta at a subapical level where Nectin-3 is co-localized with ZO1 (Fig. 6b). We did not observe such colocalization with Nectin-1, which we attribute to the poorer sensitivity of our Nectin-1 antibody staining. The colocalization between Nectin-3 and ZO1 in subapical puncta suggests that the subapical contact between IHC and PC (‘the zipper mechanism’) may also be driven by Nectin interactions.

**Figure 6.**
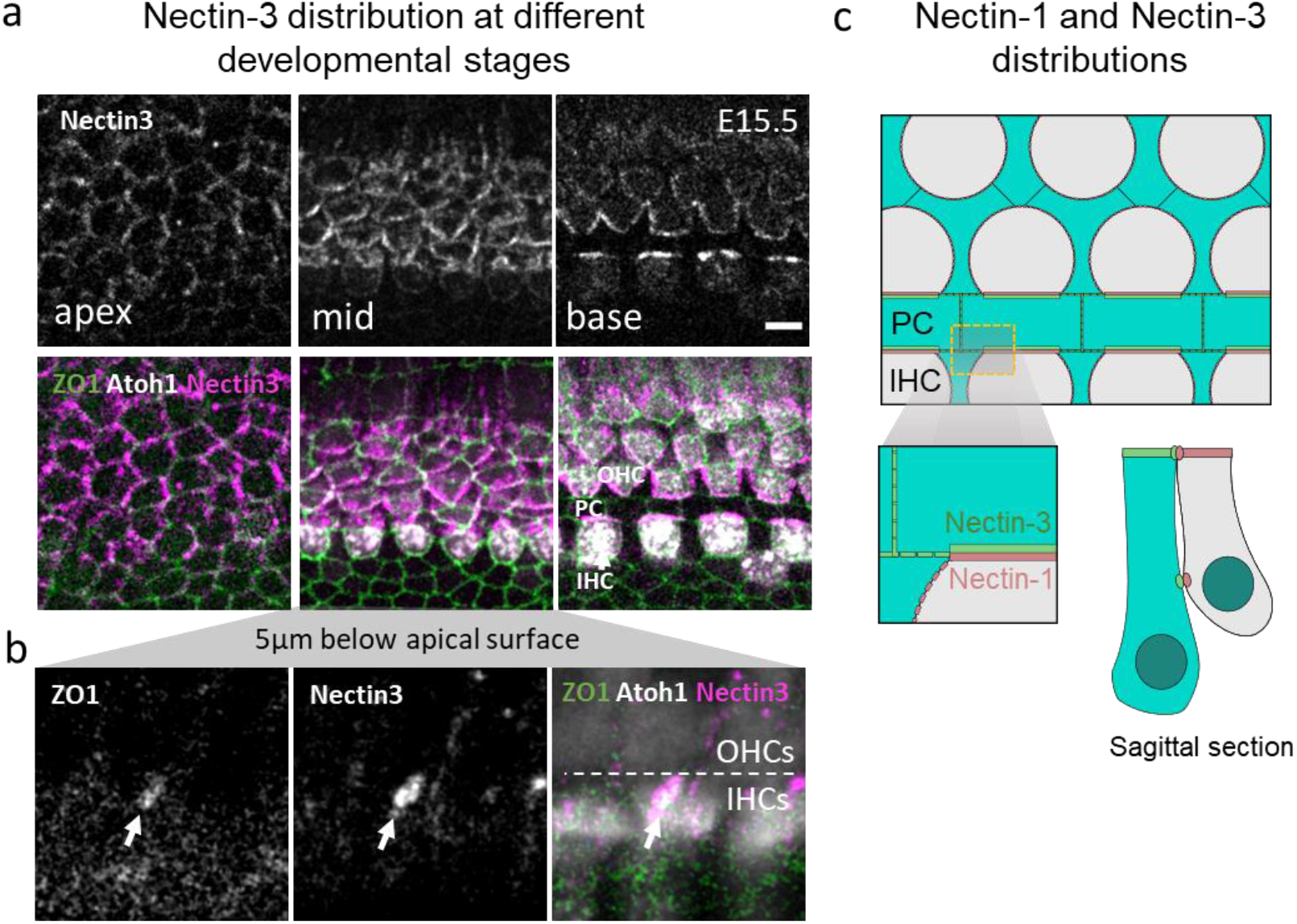
Nectin-3 accumulation at the IHC:PC boundary correlates with onset of patterning. **(a)** Antibody staining of an E15.5 cochlea with α-Nectin-3 antibody. Nectin-3 becomes localized to the IHC:PC boundary in the more developmentally advanced base region. **(b)** An image of a sub-apical plane in the mid region, showing a colocalized punctum of ZO1 with Nectin-3. Dashed line marks the PC boundary, separating between the IHC and OHC. **(c)** Schematic of the localization of Nectin-1 and Nectin-3 in the apical and sub-apical domains. Scale bar in (a): 5μm.

## Discussion

In this study we examined the process by which a single row with a precise alternating pattern of IHC and SC emerges during embryonic development. We found that an initially disordered 2-3 cell rows wide salt-and-pepper pattern is reorganized into a single row by a combination of two main processes: (i) Hopping and standard intercalations of Atoh1+ cells towards the PC row. (ii) Delaminations of out-of-line Atoh1+ cells, typically with low Atoh1 levels. These two processes resolve the two main types of defects in the initial pattern: SC:SC defects and extra out-of-line Atoh1+ cells. Further refinement of the pattern is achieved by differential adhesion between IHC, SC, and PC. This differential adhesion promotes straightening of the IHC:PC boundary and longer contact length between IHC and PC.

To elucidate the mechanism driving IHC patterning, we developed a hybrid modeling approach that captures the dynamics of regulatory circuits driven by cell-cell signaling (i.e. Notch signaling), the morphological transitions driven by mechanical forces, and the feedback between the two. More specifically, in our model, the differentiation into HC and SC affects the mechanical properties of the cells and the forces applied on them. In parallel, morphological transitions (e.g. intercalations and delaminations) and cellular morphology (e.g. apical area and boundary curvature) affect signaling by changing the identity of neighboring cells and contact length between them. Despite being based on relatively simple assumptions, the model shows that a combination of hopping intercalations and delaminations is sufficient to resolve most defects formed in the initial salt-and-pepper pattern and generate a highly organized single row of alternating IHC and SC. We note that our model does not take into account more complex regulatory relations that may affect cell fate decisions. For example, it has been shown that different Notch ligands (Dll1, Jag1, Jag2) and Notch regulators (e.g. LFng and MFng) are expressed in spatial gradients that affect Notch activity (Basch et al., 2016; Kiernan et al., 2005, 2001). Incorporating such spatial dependence of Notch activity in the model may help explain observations such as the low Atoh1 level in out-of-line cells and the decision of some of these cells to delaminate.

We have shown that a major process contributing to IHC patterning is the ‘hopping intercalation’ mechanism. This new morphological transition expands on the standard ‘tool-set’ of morphological transitions that are typically observed during epithelial morphogenesis (e.g. intercalations and delaminations) and is a result of complex sub-apical cellular dynamics and interactions. A somewhat related mechanism recently observed is that of radial intercalations, where basal cells migrate apically and open a new apical footprint at tricellular junctions (Sedzinski et al., 2016). In contrast to radial intercalations, cells performing hopping intercalations are cells that already have an existing apical surface and the general direction of the motion is lateral rather than radial.

While we do not currently know what drives the hopping intercalations in the direction of the PC, we suggest two mechanisms that are involved in this process: first, the lateral motion of HC suggests an external compression acting on HC towards the PC row, possibly driven by cell divisions in the adjacent Kolliker organ. We note that we have previously suggested a similar compression (albeit also accompanied by shear) acting on OHC driving their organization (Cohen et al., 2020). Interestingly, Atoh1+ cells detach from the basal membrane as they differentiate (Tateya et al., 2019) and become more mobile compared to SC which still remain attached. Second, the formation of sub-apical ZO1 puncta and its gradual motion towards the apical surface (i.e. the ‘zipper mechanism’) suggests that once adhesion between these cells is formed, it pulls the two cells together. The observation that these ZO1 puncta sometimes colocalize with Nectins at the contact between nascent IHC and PC suggest a molecular mechanism for the specificity of the interaction between these two cell types. Both of these mechanisms imply that forces acting at the sub-apical level can affect apical dynamics. Hence, HC patterning is a prime example for a situation where apical morphology is controlled by sub-apical dynamics, a process which is usually overlooked in most studies of epithelial morphogenesis.

The observation that some Atoh1+ cells delaminate raises the questions of what drives the decision to delaminate and what happens to the delaminating cells. We have shown that delaminating cells often have lower Atoh1 levels than their surrounding Atoh1+ cells. Recent work tracking Atoh1 activity and nuclear position in cochlear explants showed that Atoh1 level correlates with differentiation into HC. While most Atoh1+ cells move apically as they differentiate, some cells expressing lower levels of Atoh1 initially move apically, but then regress and move basally (Tateya et al., 2019). Together with our observations, this supports a picture where Atoh1 levels need to cross a certain threshold to support commitment to HC fate and for Atoh1+ cell to maintain their apical footprint. In our current imaging setup, we did not have sufficient temporal resolution to track the fate of delaminating Atoh1+ cells and thus could not determine their fate. Earlier work showed that genetic inactivation of Atoh1 in E15.5 cochleae leads to enhanced apoptosis in the prosensory epithelium (Cai et al., 2013). Hence, it is possible that the delaminating Atoh1+ positive cells apoptose. While apoptosis is a rarely observed event during normal OoC development (Cai et al., 2013), the number of delaminating cells is relatively low (see Fig. S4d) and restricted to a specific developmental stage and hence could have been overlooked.

Our model suggests that differential adhesion between different cell types helps refine the pattern by straightening the PC boundaries and adjusting the contact length between IHC and PC. It has previously been suggested that Nectin-mediated interactions drive the alternating HC pattern (Togashi et al., 2011). Based on our observations and our model, we do not believe that differential adhesion by itself could be sufficient to drive precise IHC patterning as it could not resolve many of the defects observed. Moreover, Nectin-3 knockout mice show some patterning defects but do not lead to a complete disorganization of the IHC (Fukuda et al., 2014). Hence, we suggest that differential adhesion is not a primary driving force in HC patterning, but rather a refinement mechanism.

Overall, this work highlights the important role of feedback between the regulatory level driving cellular differentiation, and the mechanical level driving morphological transitions, in establishing precise cellular patterning. Given that precise cellular patterns occur in many tissues and organs, the insights and tools developed in this work are likely to be relevant in other developmental systems.

## Acknowledgements

We would like to thank members of the Sprinzak lab and Karen Avraham for their advice and comments on this work. Atoh1-mCherry mice were a kind gift from Christine Petit, Institut Pasteur, France. ZO1-EGFP mice were a kind gift from Yasuhide Furuta and Fumio Matsuzaki from RIKEN, Japan.

## Author contributions

This scientific study was conceived and planned by R.C., S.T., O.L., and D.S. The inner ear explant dissections were performed by S.T., O.L., and S.W. Microscopy and image analysis were performed by R.C., S.T., and S.K., Modeling was performed by R.C. Manuscript was written by R.C., S.T., O.L., and D.S.

## Tables

**Table S1.** Model parameters. The parameters used in each one of the models. See methods for definitions of parameters.

| Simulation | Lateral inhibition parameters | Vertex model parameters |
| --- | --- | --- |
| L.I. only model<br>(Figure 1d) | $\beta_N = \beta_D = 3.9, \beta_R = 194,$<br>$\gamma_R = 60, k_t^{-1} = 0.2,$<br>$m = l = 3$ | |
| hybrid 1 model<br>(Figure 3d) | Same as L.I. only model | $A_0 = \frac{4\pi^2}{N_c}, \gamma_0 = 0.05, \Gamma_0 = 0.1,$<br>$\zeta_0 = 1, \sigma_0 = 1, \kappa = 12, D = 0.7, q = 3,$<br>$\alpha_{HC} = 5, \alpha_{other} = 1, \Gamma_{HC} = 2\Gamma_0, \Gamma_{other} = \Gamma_0$ |
| hybrid 2 model<br>(Figure 4e) | Same as L.I. only model | Same as hybrid 1 model + delamination of out-of-line HC |
| hybrid 3 model<br>(Figure 5d) | Same as L.I. only model | Same as hybrid 2 model + adhesion rules:<br>$\gamma_{HC:HC} = \gamma_{SC:PC} = 10\gamma_0, \gamma_{HC:PC} = 7\gamma_0, \gamma_{other} = \gamma_0$ |

**Table S2.**
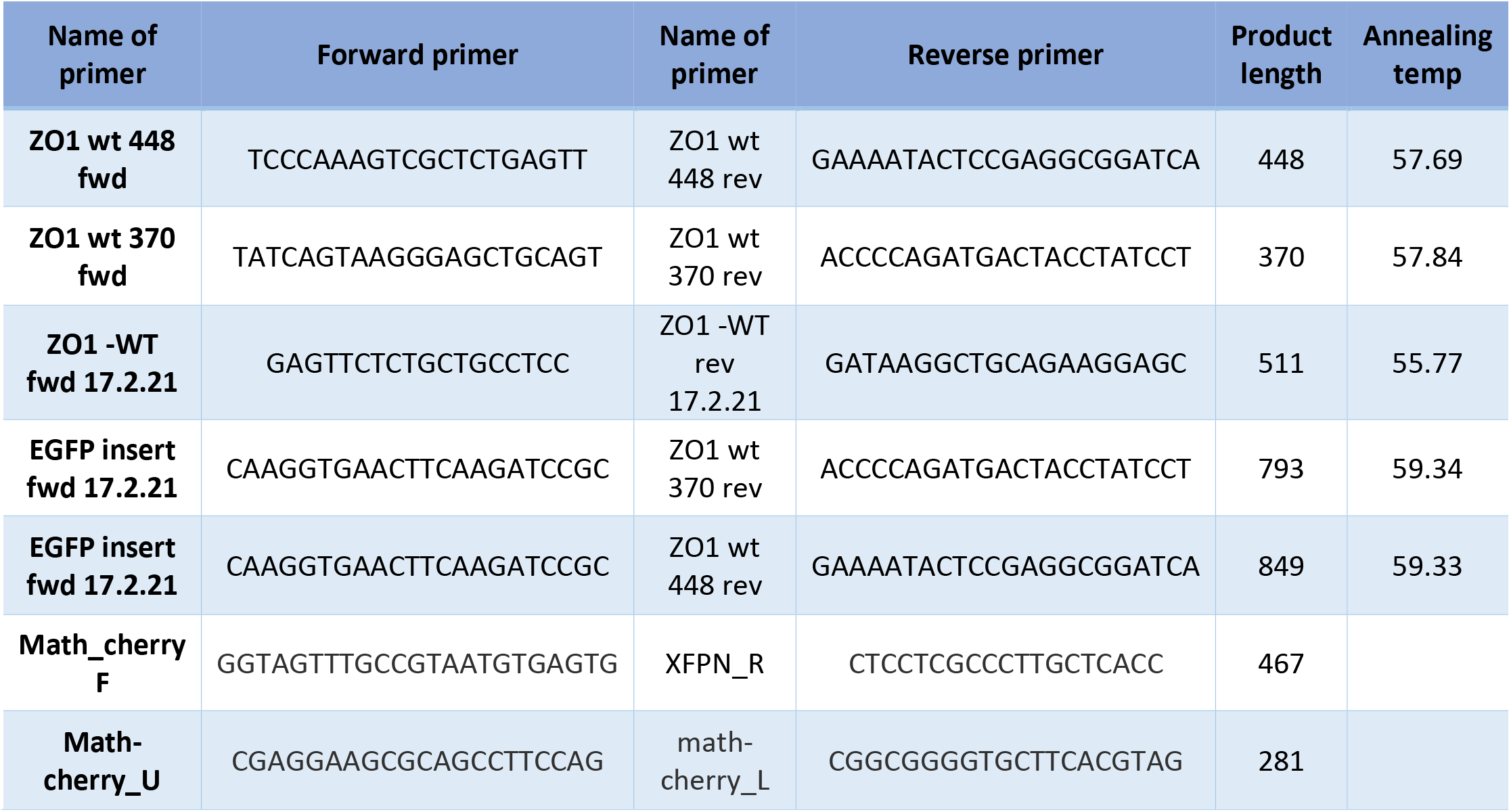
Primer list. A list of the primers used for genotyping.

## Materials and methods

### Mice

Atoh1-mCherry mice were a gift from Christine Petit, Institute Pasteur, and were maintained on a C57bl/6J background. Rosa26-ZO1-EGFP mice were a gift from Yasuhide Furuta and Fumio Matsuzaki from RIKEN Laboratory13 (accession no. CDB0260K) and maintained on a C57BL/6 background. All animal procedures were approved by the Animal Care and Use Committee at Tel Aviv University (04-20-003). Genotyping was performed using the KAPA HotStart Mouse Genotyping Kit (Sigma, KK7352) using primers listed in Supplementary Table 2. For timed mating, one or two females 8-15 weeks old were paired with appropriate male for overnight mating. Mice were separated the next morning.

### Immunohistochemistry

Neonatal mice at P0 were sacrificed by decapitation, and adult pregnant mice were sacrificed by CO2 inhalation. The inner ears were dissected from P0 pups and E15.5 embryos directly into cold PBS and fixed in 4% paraformaldehyde (Electron Microscopy Sciences, cat: 15710) for 2 hrs at room temperature. For whole-mount imaging, sensory epithelia were exposed and separated from the inner ear. Next, samples were incubated in 10% normal Donkey serum (Sigma, cat: D9663) with 0.2% Triton-X (Sigma, cat: T-8787) for 2 hrs at room temperature. Samples were then incubated overnight at 4°C in the appropriate primary antibody diluted in antibody diluent buffer (GBI labs cat: E09-300). Following 3 washes in PBS, samples were incubated with secondary antibodies for 2 hrs at room temperature. Stained samples were mounted on Cover glass 24×60mm thick. #1(0.13-0.17mm) slides (Bar-Naor Ltd. cat: BN1052441C) using a fluorescent mounting medium (GBI, cat: E18-18). Image acquisition was done with a ZEISS LSM 880 with Airyscan microscope (Zeiss). List of antibodies and dyes: anti MyoVIIa 1:250 (Proteus Biosciences, cat: 25-6790), anti Nectin-3 1:250 (Genetex, cat: GTX16913), anti Nectin-1 1:250 (MBL, cat: D146-3), Alexa fluor 647 donkey anti rat 1:500 (Jackson laboratories, cat: 712-605-150), Cy™3 AffiniPure Goat Anti-Mouse IgG (H+L) 1:500 (Jackson laboratories, cat: 115-165-062), Alexa Fluor™ 633 Phalloidin 1:50 (Thermo, cat: A22284).

### Organ of Corti explants

Cochlear dissection for ex-vivo culture was performed as previously described (Cohen et al). Briefly, cochleae were dissected in cold PBS and placed in a Cellvis 4-chamber glass bottom dish (D35C4-20-1.5-N) inside 100 ul of Matrigel Phenol Red Free solution (In vitro technologies, cat: FAL356237) diluted 1:15 in DMEM. After Matrigel solidified, medium was added, and samples were transferred to the imaging chamber. DAPT treatments: DAPT was diluted according to manufacturer’s instructions in DMSO. The diluted solution was then added to the culture dish at the beginning of the movie to reach a final DAPT concentration of 1 μM. Control dishes contained the same amount of DMSO without DAPT.

### Microscopy details

Cochleae were imaged using Zeiss LSM 880 confocal microscope equipped with an Airyscan detector using 488nm laser for GFP and 561nm laser for mCherry and 633nm for Alexa Fluor 647. For fixed samples we used Plan-apochromat 63x oil-immersion objective with NA=1.4. For live imaging we used C-apochromat 40x water-immersion objective with NA=1.2. The microscope was equipped with a 37 °C temperature-controlled chamber and a CO2 regulator providing 5% CO2. The equipment was controlled using Zeiss software – “Zen black”.

### Laser ablation system

Laser ablation experiments were performed using Rapp OptoElectronic UGA-42 Caliburn system equipped with a 355nm UV short pulse laser. The ablation system was integrated to the microscope’s hardware and software and controlled using Rapp software SysCon. In ablation experiments, cells were hit with 500 short consecutive pulses over 0.5 seconds with a laser power of 4%.

### Image analysis

Images of the OoC were taken as a sequence of focal planes (z-stack) spanning a depth of 20-30*μm* with a pixel size of 0.1μm and separation of 0.7μm between z-stacks ([0.1, 0.1, 0.7]μm/voxel). Movies are a sequence of such images with a frame rate 1/15min. Laser ablation movies were taken with a smaller field of view and with a faster frame rate of 1/2min. Data processing was performed off-line using custom built codes in Matlab (MATLAB R2020b, the MathWorks Inc.) and using the open-source software ImageJ. The presented images and movies were obtained from a custom-made surface projection algorithm in Matlab (see “Surface projection” section). Images of sagittal planes were produced by using ImageJ’s Orthogonal Views function.

Segmentation of the cellular boundaries was done using the ZO1 channel in the surface projection. The images were pre-adjusted for segmentation by using gaussian blur filter and h-minima transform. Then, a watershed algorithm was applied to get a binary image of the cellular boundaries. Any defects in the segmentation were manually fixed using a custom-made code.

To get data of individual cells we used the labeled segmentation image and the Atoh1 image. HC, SC and PC were detected and marked manually according to Atoh1 expression, cell position and cellular morphology. For each cell, the following data was trivially extracted: apical surface area, cell circumference, Atoh1 expression and neighboring cells.

### Surface projection

Our data is volumetric 3D stacks of sections of the OoC, that contain the apical surface as well as sub-apical surfaces. Since ZO1 is concentrated in tight junctions, we used maximum intensity projection (MIP) to get a 2D projection of the cellular boundaries at the apical surface. For Atoh1 however, MIP creates a projection of the entire cellular volume of HC, not only from the apical surface. Therefore, using MIP for a two channel ZO1-Atoh1 image makes it hard to relate Atoh1 expression with a specific cell observed at the apical surface. For that reason, we developed a surface projection algorithm that presents a 2D projection of the apical surface for both ZO1 and Atoh1 as well as reduces off-apical noise.

The first step in the algorithm was to create a stack that roughly represents the apical surface. As ZO1 is brightest at the apical cellular boundaries, we use strong gaussian blurring (gaussian sigma ~3*μm*) to get a smooth stack that roughly represents the apical surface (surface stack). The second step is to refine the surface stack and get a 3D mask that represents the apical surface more accurately (surface mask). For that, we started with an empty stack with identical dimensions to the surface stack. Then, for each xy-position in the surface stack, we find the z-position with the highest ZO1 signal and mark the corresponding voxel in the surface mask as 1. At this point the surface mask contains 1 at the voxels that represent the apical surface or 0 otherwise. To make the surface mask smoother we apply a weak gaussian blur filter (gaussian sigma ~0.2*μm*). We then multiply the original ZO1 stack and the Atoh1 stack with the surface mask (voxel to voxel multiplication). These new stacks contain the signal from the apical surface alone. Finally, we take MIP of these new stacks to get a 2D projection of the apical surface.

### Change in apical area after hopping intercalation

To measure the change in apical area of cells that perform hopping intercalations we measured the apical area of each hopping cell before and after hopping. The first measurement was done one frame before a new apical surface was first observed (~15min before) and the second measurement was done 1 hour after the two apical surfaces were merged, to allow enough time for relaxation. As a control, we measured the apical area of non-hopping IHC neighbors at the same timepoints. This average change for hopping and non-hopping cells is presented in Fig. 2e.

### Circularity of IHC

To assess the roundness of IHC at different developmental stages we used the circularity measure. Circularity is defined as 4*πA/L^2^* where *A* is the cell apical area and *L* is the cell circumference. The value range for circularity is 0-1, where values closer to 1 indicate that the shape is closer to a perfect circle. We analyzed images from E15.5 and E17.5 cochleae, indicating early and advanced timepoints, respectively. For each image we measured the circularity of 10 randomly chosen IHC and calculated the average value for that image. We then averaged the circularity values of different images from E15.5 and E17.5 cochlea separately, as presented in Fig. 3c.

### Number and Atoh1 levels of delaminating Atoh1+ cells

To measure the Atoh1 expression of the cells we used the mCherry channel in the surface projection and the segmented image. For each cell, we defined the Atoh1 level as the total mCherry intensity of the cell divided by the area of the cell. The intensity of mCherry can vary between different experiments due to differences between mice. Therefore, we defined a normalized Atoh1 level as the Atoh1 level of the cell divided by the average Atoh1 level of the 10 closest neighboring Atoh1+ cells. This normalized level indicates the relative intensity of the cell with respect to other Atoh1+ cells. For each delaminating Atoh1+ cell, the intensity measurement of the cell and its Atoh1+ neighbors was performed one frame before the delamination. The averaged normalized intensity of delaminating cells and their neighbors is presented in Fig. 4b.

To link between the decision of Atoh1+ cells to delaminate and their position, we counted the number of delaminating Atoh1+ cells that touch/do not touch the PC row at the time of delamination. Additionally, we counted the number non-delaminating Atoh1+ cells that touch/do not touch the PC row at the end of each movie. PC were identified by their rectangular morphology. If the analysis was done at a stage where cellular morphology was not yet differentiated, the PC were identifying at a later timepoint in the movie, where morphological differences are clear, and then traced back to the analyzed timepoint. The percentage of delaminating/non-delaminating Atoh1+ cells that touch/do not touch the PC row is presented in Fig. 4c.

Finally, to link between Atoh1+ cells position and their Atoh1 level, we calculated the normalized Atoh1 level for each cell that touch\do not touch the PC row at the initial point of each movie. The averaged intensity for touching\non-touching cells is presented in Fig. 4d.

### Effect of Notch inhibition on Atoh1+ cells

To investigate the effect of Notch inhibition in early developmental stages, we imaged E15.5 cochlear explants for 3 hour, then treated them with 1μM of DAPT and continued imaging for 20 hours. We manually counted the number of Atoh1+ cells in the IHC region at the last timepoint before DAPT treatment and 20 hours after treatment, as well as the number of delaminations of Atoh1+ cells during the movies. For each movie we calculated the ratio between the final and initial number of Atoh1+ cells and the number of Atoh1+ cells per 100μm. As a control, we repeated the same measurements for E15.5 cochlear explants treated with DMSO. The averaged data of the DAPT and DMSO movies is presented in Fig. S4b-d.

### Morphological measurements of PC boundary

To estimate morphological parameters of the PC row at different developmental stages we used images from E15.5 and E17.5 cochleae. To estimate the straightness of the PC row we manually marked the center of each of the PC in each image and used a modified version of the reduced chi-squared statistic given by:

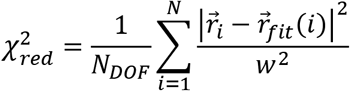

where *N* is the number of PC, 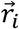 is the position of cell *i*, 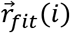 is the closest point to 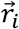 on the best fitted line passing through all marked positions, *w* is the weight and *N_DOF_* = *N* – 2 is the degrees of freedom. We chose the weight to be the average horizontal length of a PC given by 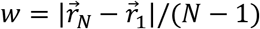. The average 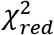 of the PC row in different developmental stages is presented in Fig. 5b.

To measure the contact length between IHC/SC and the PC row we segmented the IHC:PC and SC:PC boundaries in E15.5 cochleae. For each IHC/SC that touches the PC row, we measured the total length of its interface with the PC row (the cell can have boundaries with more than one PC). The average PC row contact length for IHC and SC is presented in Fig. 5c.

### PC ablation experiments analysis

To determine the effect of mechanical perturbations in the PC row at different developmental stages, we ablated PC in the base and mid regions of E17.5 cochleae. In each experiment, a single PC was ablated using a UV laser pulse directed at the center of the apical surface of the cell (see “Laser ablation system” for more details). Immediately after the ablation, a small region around the ablated PC was imaged for 30 minutes with a 1-minute interval, during which the continuity of the PC row was either broken or repaired by the two neighboring PC. The percentage of experiments that resulted in PC row repair/break in the mid/base regions is presented in Fig. 5j. We note that some of the ablation experiments were performed in different positions on the same cochlea, separated by at least 10 cells diameter. Overall, 13 cochleae were used for PC ablation experiments.

### Lateral inhibition model

We simulate lateral inhibition patterning based on our previously described cell-cell contact dependent lateral inhibition model (Shaya et al., 2017) that incorporates the assumption that signaling strength between two cells is proportional to the contact area between the cells. This is a simplified model that assumes only one kind of receiver – Notch, and one kind of ligand – Delta. Furthermore, for simplicity, we considered a model that does not include cis inhibition between receptors and ligands. Here, we present a short summary of the model.

Assume that cell *i* and cell *j* are neighbors and share a boundary with a contact length of *l_ij_*. We denote Notch concentration in cell *i* at the boundary with cell *j* as *n_ij_*. Similarly, *d_ij_* is Delta concentration in cell *i* at the boundary with cell *j*. The signaling scheme is simplified as follows: Notch on cell *i* interacts with Delta on cell *j* with an association rate of 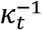, and produces a signal in cell *i* proportional to *n_ij_d_ji_l_ij_*. The total repressive signal received by cell *i*, *R_i_*, is proportional to ∑*_j_ n_ij_d_ji_l_ij_*, with the sum being over all the neighboring cells to cell *i*. Therefore, the unitless dynamical equations for Notch concentration, Delta concentration and the repressor expression can be written as:

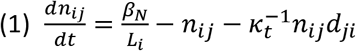

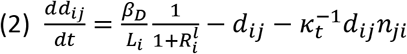

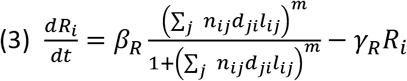

where *β_N_,β_D_,β_R_* are the production rates of Notch, Delta and the repressor respectively, *γ_R_* is the degradation rate of the repressor, *L_i_* is the perimeter of cell *i*, and *l*, *m* are the Hill coefficients for the repressor and total signal respectively. For complete details of the lateral inhibition model see (Shaya et al., 2017).

### Simulations of lateral inhibition model

To run the lateral inhibition model, we need to have a defined tissue geometry and adjacency information of neighboring cells. The geometry of the tissue is defined on the basis of the vertex model, such that each cell is represented by a polygon in a connected network of cells (see 2D vertex model section). This representation allows for the definition of neighboring cells and boundary lengths.

As differentiation of IHC occurs in a narrow region of cells, we define the signaling domain for lateral inhibition as a region of cells spanning 2-3 rows. As an initial state, cells in the signaling domain start with randomly distributed basal levels of Notch and Delta, and no repressor. Cells in the medial side of the signaling domain are kept with a constant low level of Notch ligand but without Notch receptors, to account for the low levels of Jag1 observed in the Kolliker organ (Basch et al., 2016). All other cells do not express Notch receptors or ligands.. Given this defined initial state, the model is advanced in time in an iterative manner, by changing the expression values according to equations (1)-(3).

Finally, we want to consider the diffusion of Notch and its ligand over the cell membrane. While for each cell the model changes the concentrations of Notch and its ligand on individual cellular boundaries, we assume that these concentrations diffuse across the circumference of the cell in a faster timescale relative to the simulation timesteps. To implement this, we homogeneously redistribute the total expression of Notch and its ligand over the cellular boundaries of the cell, between each two timesteps in the simulation.

Simulation codes for the lateral inhibition model can be found in the public repository https://github.com/Roie-Cohen/mechano-signaling-inner-ear-model.git.

### 2D vertex model

2D vertex models are widely used for simulating the dynamics of epithelial tissues (Farhadifar et al., 2007). In 2D vertex models, cells in the tissue are represented by a connected network of polygons, with each side of the polygon representing a cellular boundary between neighboring cells and each vertex representing a tri-cellular junction. In this simplistic description of epithelial tissues cells can only have straight cellular boundaries with their neighbors, however, in cases where boundary curvature is significant this description is not sufficient. Since HC are significantly rounder than other cell types in the OoC, we solve this problem by introducing additional virtual vertices on each cellular boundary, effectively allowing the boundaries to curve (Frost et al., 1988; Tamaki et al., 2012).

With the geometry of the tissue defined, mechanical energy can be assigned for each cell, cellular boundary, or any other local or external (conservative) force that might act on the cells. The total mechanical energy of the system is the sum of all these terms (Fig. S3a). The dynamics of the tissue is determined according to the gradient descent method, by iteratively repositioning all vertices in the direction that minimizes the total mechanical energy of the system (Farhadifar et al., 2007). To implement a hybrid model that combines cellular dynamics with lateral inhibition, we run both the vertex model and the lateral inhibition model in parallel, such that both models are advanced in time simultaneously (for further details see “Simulations of hybrid models”).

Simulation codes for all hybrid models can be found in the public repository https://github.com/Roie-Cohen/mechano-signaling-inner-ear-model.git.

### Defining cell types

The forces and interactions between cells can be different for different cell types. Therefore, to appropriately define the energy terms in the model, we assign each cell as either HC, SC, PC or a general type cell. HC and SC naturally emerge from the signaling domain in which lateral inhibition is applied. Each cell in the signaling domain whose ligand expression passes a certain threshold is set as a HC, while the other cells in the domain are considered as SC. Each cell that borders the signaling domain on its lateral side is set as a PC. All other cells outside the signaling domain are defined as general type cells.

### Mechanical energy function

The total mechanical energy of the system can be separated into terms resulting from (i) the internal mechanical properties of the cells, (ii) interaction between cells and (iii) external forces:

#### (i) Internal energy terms

The internal terms for the energy function account for the compressibility and contractility of the cells. The compressibility accounts for energy needed to compress the cell, and is given by:

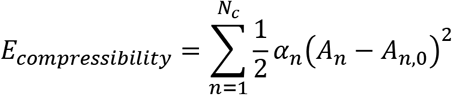

where *α_n_* is the incompressibility of cell *n, A_n_* is the area of the cell, *A_n,0_* is the preferred area of the cell and *N_c_* is the number of cells in the tissue.

The contractility term accounts for the contracting actomyosin cables at the perimeter of the cell, and is given by:

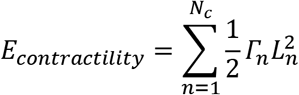

where *Γ_n_* is the contractility of the cell and *L_n_* is the circumference of the cell.

#### (ii) Interaction terms

The interaction terms in the energy function account for the adhesion and repulsion between cells. The adhesion term is given by:

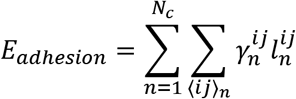

where 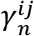 and 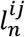 are the tension and length of the boundary of the cell that lies between vertices *i* and *j*, respectively. The second sum is done over all the boundaries of cell *n*.

The repulsion between cells acts only between HC and corresponds to the physical contact between HC at the HC nuclei level. The simplest way to model such repulsion is using a hard-sphere potential, however, such a term is non-continuous and does not have a well-defined derivative. To capture a similar behavior with a continuous term, we use the following:

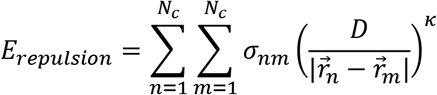

where *σ_nm_* is the repulsion strength between cells *n* and *m, κ* is the repulsion exponent, *D* is the effective diameter of the HC nuclei and 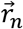, 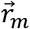 are the center of mass (COM) positions of the cells. As the repulsion act only between HC we have:

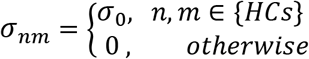

By setting the repulsion exponent κ to a high value, we create a term with a very strong repulsion for cells with touching nuclei 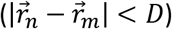 and a very weak repulsion for cells with non-touching nuclei 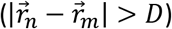. Since there should be no long-range repulsion between cells we set a cutoff distance, such that this term vanishes for 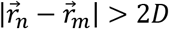. Notice that since κ is large, the discontinuity we create is negligible.

#### (iii) External forces

We model the lateral compression on HC as a linear force that act on HC in the direction of the PC row. The energy associated with the compression force is then:

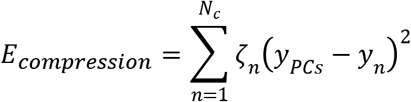

where ζ_n_ is the compression force, *y_PCs_* is the average *y* position of the COM of PC and *y_n_* is the *y* position of the COM of cell *n*. We additionally assume that this compression force does not act on cells that are already close enough to the PC. To implement this, we add an additional cutoff rule: once a cell is one cell diameter apart or less from the PC, the force vanishes. Finally, as lateral compression act only on HC we have:

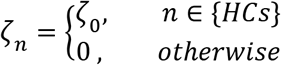

#### (iv) Unitless parameters

For convenience, we choose to work with unitless parameters. The energy function is normalized by *α*_0_*Ā*^2^, where *α*_0_ is the incompressibility of SC and *Ā* is the average area of the cells. This accounts for the following transformation:

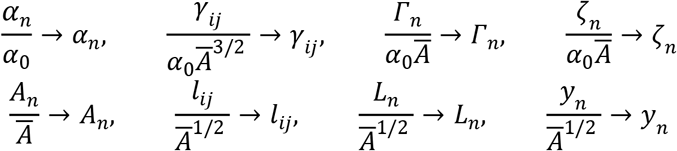

### Modeling hopping intercalation

To model hopping intercalation in the vertex model we define a hopping event as a process where a cell opens a new apical surface that expands, and finally merges back into one apical surface. We define three conditions for the initiation of a hopping event: (a) the hopping cell must be a HC, (b) the hopping cell and the PC row must be separated by one cellular boundary, and (c) the area of the hopping cell must be 90% or less than its preferred area. The first and second conditions restrict hopping events to HC that hop to the PC row, as observed in our movies. The third condition sets an internal pressure threshold, effectively defining an energy barrier that the cell must overcome to hop over the boundary.

Once a hopping event is initiated, a new apical surface is opened at a tri-cellular junction at the PC row. This new apical surface is modeled as a new cell that is linked to the original hopping cell. The new cell initially adopts all mechanical properties and Notch/ligand expression levels of the original cell. The total expression of Notch/ligand is homogeneously redistributed over the boundaries of the two cells, between each two timepoints of the simulation. The mechanical linkage between the cells is expressed by the following modification to the compressibility energy term:

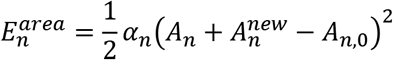

where *A_n_* and 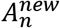 are the areas of the original and new apical surfaces of cell *n*. This modification captures the fact that the two cells share the same cellular volume. Additionally, as the expansion of the new apical surface requires additional energy (that pushes the neighboring cells), we add an expansion force to the new apical surface in the form of:

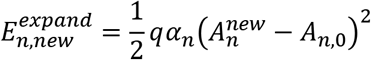

where *q* is a positive expansion factor that determines the strength of the expansion force. This force is possibly originated by the cellular body, pushing the new apical surface up.

### Initial and boundary conditions

The initial configuration for the lateral inhibition model and for the hybrid 1 and hybrid 2 models is a disordered lattice of cells. To generate disordered cell lattices we create 10×25 hexagonal lattices and assign random values for boundary tensions (*γ_ij_*) and preferable areas (*A*_0,*n*_). We then run the vertex model for each lattice and reassign random values for the parameters every few timesteps. Once we have a disordered lattice, we define the signaling domain and assign cell types (as described in “Defining cell types”). For simplicity, in all simulations we use periodic boundary conditions at the left-right and top-bottom edges of the cell lattice (boundary conditions on a torus).

### Simulations of hybrid models

In this work we present three hybrid models that combine our lateral inhibition model with different 2D vertex models. The progression of each hybrid model is done by iteratively changing the Notch/ligand expressions of the cells and the cellular configuration of the lattice, at the same time. In parallel, we allow both intercalations and delaminations to take place for each boundary smaller than a defined threshold length and for each cell smaller than a defined threshold area, respectively. In addition, we allow HC to perform hopping intercalations into the PC row (see details in “Modeling hopping intercalation”).

In the hybrid 1 model, as presented in Fig. 3d, we assume a lateral compression force that acts on HC and local repulsion force that acts between HC (see “Mechanical energy function” for details). The lateral compression pushes the HC towards the PC row and drives either standard intercalations or hopping intercalations. In addition, as HC have a round morphology (Fig. 3c) and lower deformability relative to SC (Cohen et al., 2020), we assign HC with higher contractility (*Γ_n_*) and higher incompressibility (*α_n_*) compared to other cell types.

As described in the main text, the hybrid 1 model does not account for the observation of delaminating Atoh1+ cells. To include this observation, we modify the hybrid 1 model by adding a delamination rule: each HC that does not touch the PC row for a long enough period (defined by a threshold), delaminates. Delamination of a specific cell is forced by setting the preferable area to zero and increasing the contractility to a very high value, causing the cell to rapidly shrink below the threshold area for delamination. Adding this simple delamination rule removes all out-of-line IHC defects in the final pattern, as shown in the simulation in Fig. 4e. We refer to this modified model as hybrid 2 model.

While the hybrid 2 model captures both hopping intercalations behavior and the reduction in the number of defects, it does not capture the longer length of the IHC:PC boundaries and the high straightness of the PC row. To include these morphological aspects, we added a second stage to the model (hybrid 3 model), that starts at the final timepoint of hybrid 2 model simulations. In this stage we increase the tensions of the HC:PC and SC:PC boundaries such that *γ_IHC:PC_* > *γ_SC:PC_* > *γ*_0_, where *γ*_0_ is the tension in SC:SC boundaries. Additionally, we increase the tension in the HC:HC boundaries. We note that while we introduce the adhesion rules as a second stage to the model, the two stages likely overlap.

The values for all parameters can be found in Supplementary Table 1.

### Simulations of single PC laser ablation

To investigate the effect of the adhesion rules, PC ablation simulations started at the end of either hybrid 2 model (without adhesion rules) or hybrid 3 model (with adhesion rules). In each simulation, a single PC was “ablated” by setting the preferable area to zero and increasing the contractility to a very high value, causing the cell to rapidly shrink and delaminate. In each set of simulations (with\without adhesion rules), we counted the number of times the PC row repaired or broke after the ablation. The data is presented in Fig. 5h.

### Model results analysis

The measurements of circularity, change in area after hopping intercalation, reduced chi-squared of the PC row, contact lengths of IHC:PC and SC:PC interface and PC row break rate after PC ablation were performed exactly as in the experiments.

Number of defects was counted at the final timepoint of the simulations of the hybrid 1 model (Fig. 3d). Out-of-line HC defect is defined as any HC that does not have a boundary with a PC. SC:SC defect is defined as an occurrence of two adjacent SC that both have a boundary with PC.

### Statistical analyses

Statistical analyses were performed using Prism 8 software (GraphPad, San Diego, CA). The details of the statistical tests used are found in the figure legends. To test data for normality of distribution, the Anderson-Darling, D’Agostino & Pearson, Shapiro–Wilk and Kolmogorov–Smirnov tests were used. If a dataset did not pass the normality threshold in at least one of these tests, the Mann-Whitney test was used. If data passed normality tests but did not pass the F test for comparing variances, Welch’s t-test was used. If variances were not significantly different, Student’s t-test was used. For comparison of discrete categorical data, the Chi-square test was used.

## Data availability

Data supporting the findings of this manuscript are available from the corresponding author upon reasonable request.

## Code availability

Matlab codes for lattice generation and running the simulations were uploaded to the public repository https://github.com/Roie-Cohen/mechano-signaling-inner-ear-model.git.

## Supplementary Figures

**Figure S1.**
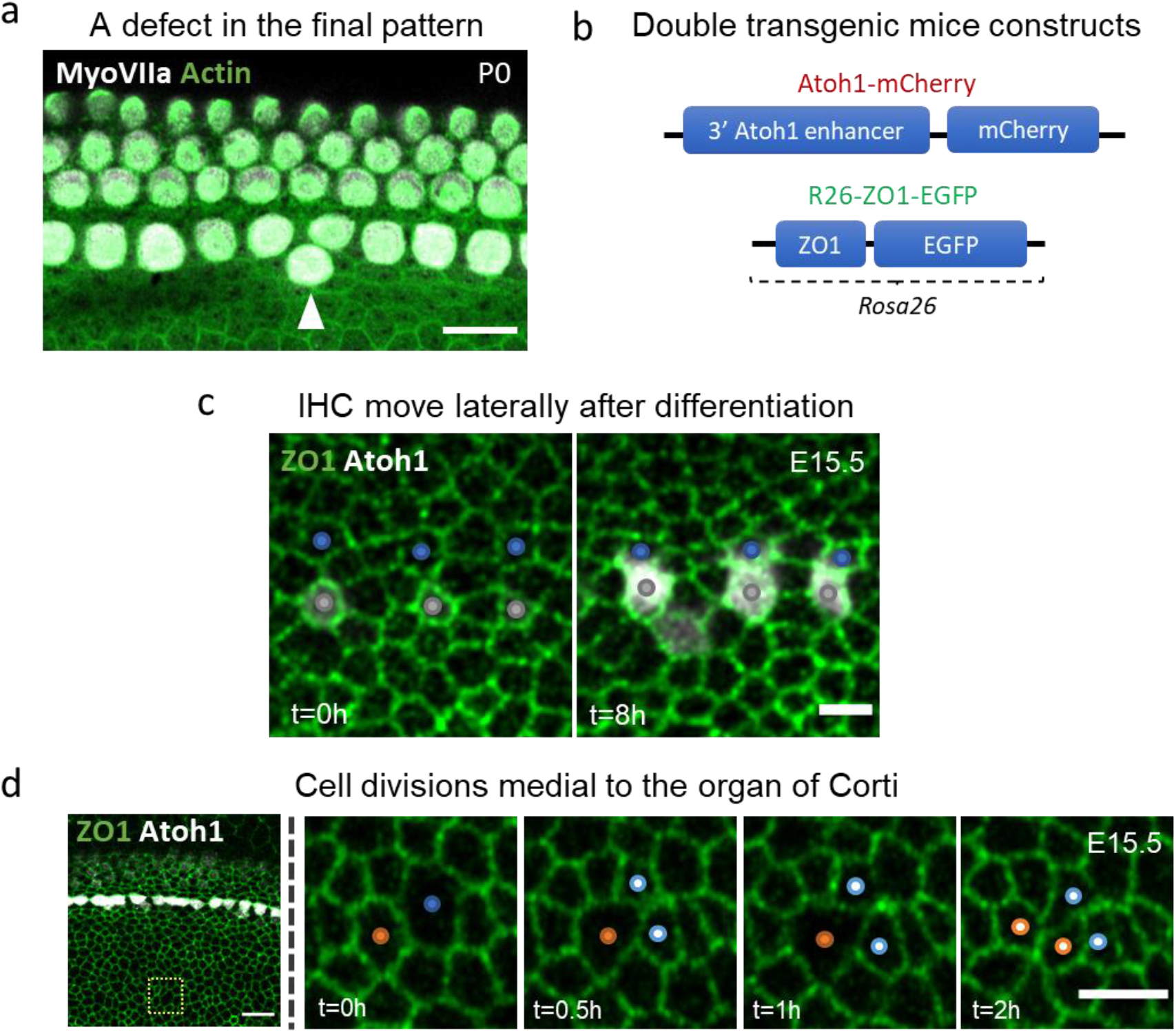
Dynamic movements of Atoh1+ cells during IHC formation. **(a)** An image of a P0 cochlea, stained for MyoVIIa (marks HC, white) and actin (green). Arrowhead marks a defect in the final pattern, in the form of an extra out-of-line IHC. **(b)** Genomic architecture of reporter systems in double transgenic mice used for live imaging. **(c)** Images at two time points from a movie of an E15.5 cochlear explant at the apex region showing relative lateral positions of cells before and after IHC differentiation. After differentiation, the Atoh1+ cells (gray dots) move laterally toward the future PC (blue dots). **(d)** A filmstrip from an E15.5 cochlear explant from a region medial to the IHC (dashed box in left panel) showing dividing cells (blue and red dots) and their respective daughter cells (light blue and light red dots) (Movie S4). Scale bars: 10μm for (a) and left panel in (d), 5μm for (c) and filmstrip in (d).

**Figure S2.**
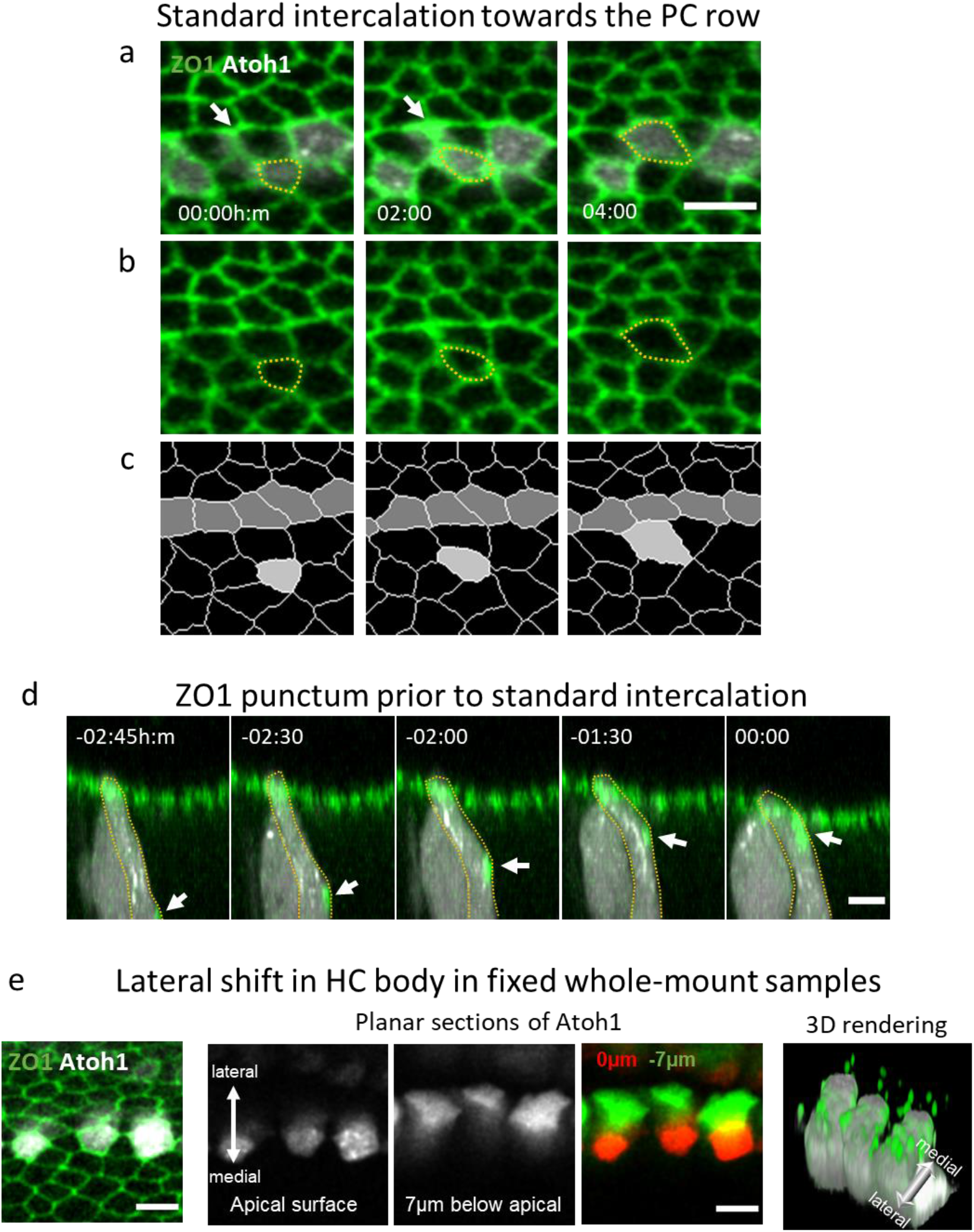
Atoh1+ cells also perform standard intercalations towards the PC row. **(a-c)** A filmstrip from a movie of an E15.5 cochlear explant showing an apical view of a standard intercalation event where (a) shows both Atoh1 and ZO1 markers, (b) shows only the ZO1 marker, and (c) shows a segmentation of (b) highlighting intercalating cell (light gray) and PC (dark gray) (Movie S5). Arrow in (a) shows the destination of the intercalating Atoh1+ cell. **(d)** A cross section of the filmstrip in (a-c) showing a ZO1 punctum propagating towards the apical surface. Timestamps show time with respect to the intercalation event (at 00:00h:m). **(e)** An image of a fixed E15.5 cochlea confirming the observed lateral shift of HC bodies in cochlear explant experiments. Scale bars: 5μm.

**Figure S3.**
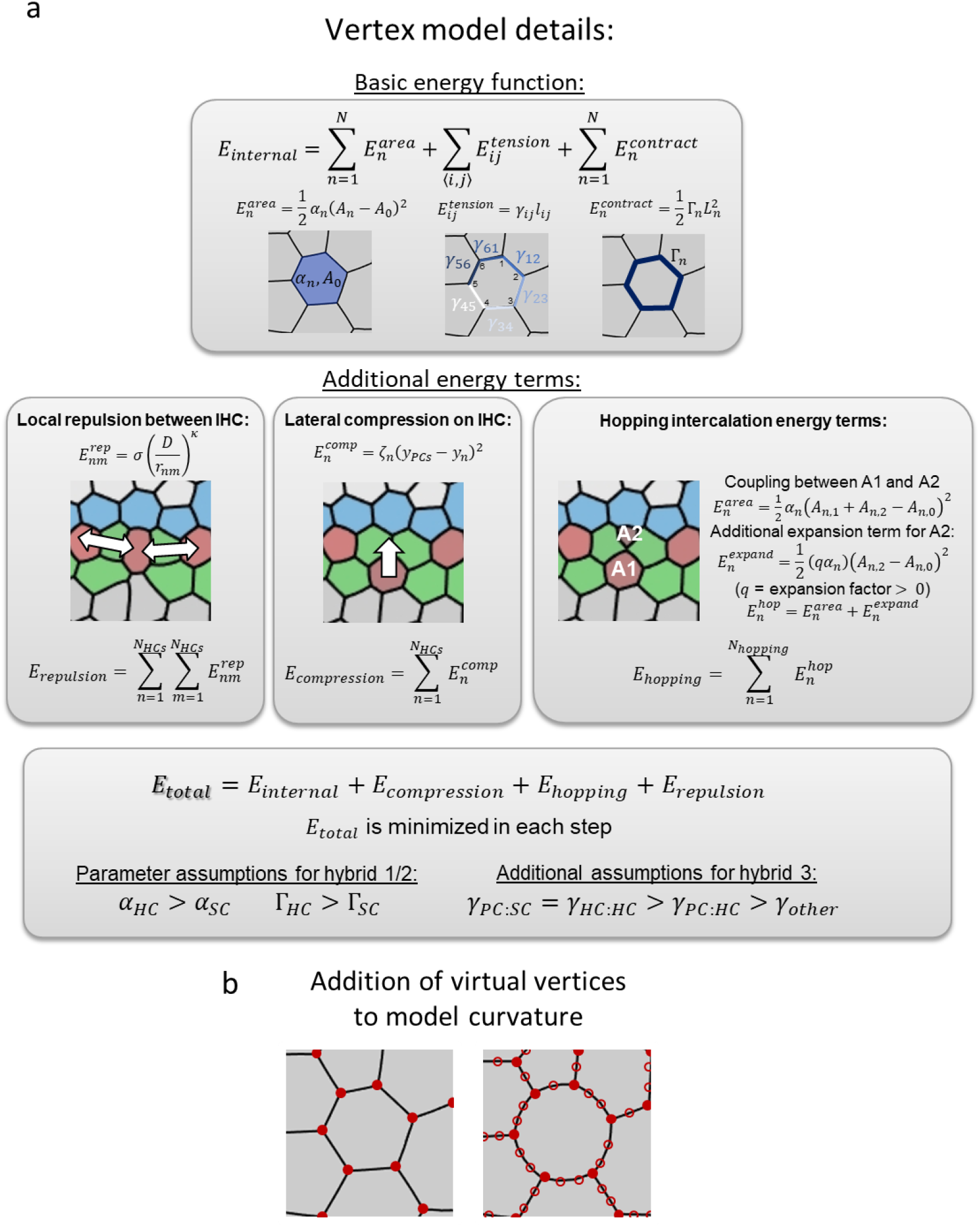
Schematic description of the terms in the 2D Vertex model. **(a)** Schematic description of the 2D vertex models used. First row shows the standard energy terms associated with cell area (incompressibility), boundary length (line tension), and circumference (contractility). The second row shows the additional energy terms associated with local repulsion between IHC, lateral compression on IHC, and hopping intercalation. In each step of the simulation the total energy is minimized. Assumptions regarding the parameters in the different models are shown in the bottom row. See full details in methods. **(b)** Schematic of virtual vertices. In all models, in addition to the tricellular vertices (filled circles), virtual vertices (empty circles) were added to each edge.

**Figure S4.**
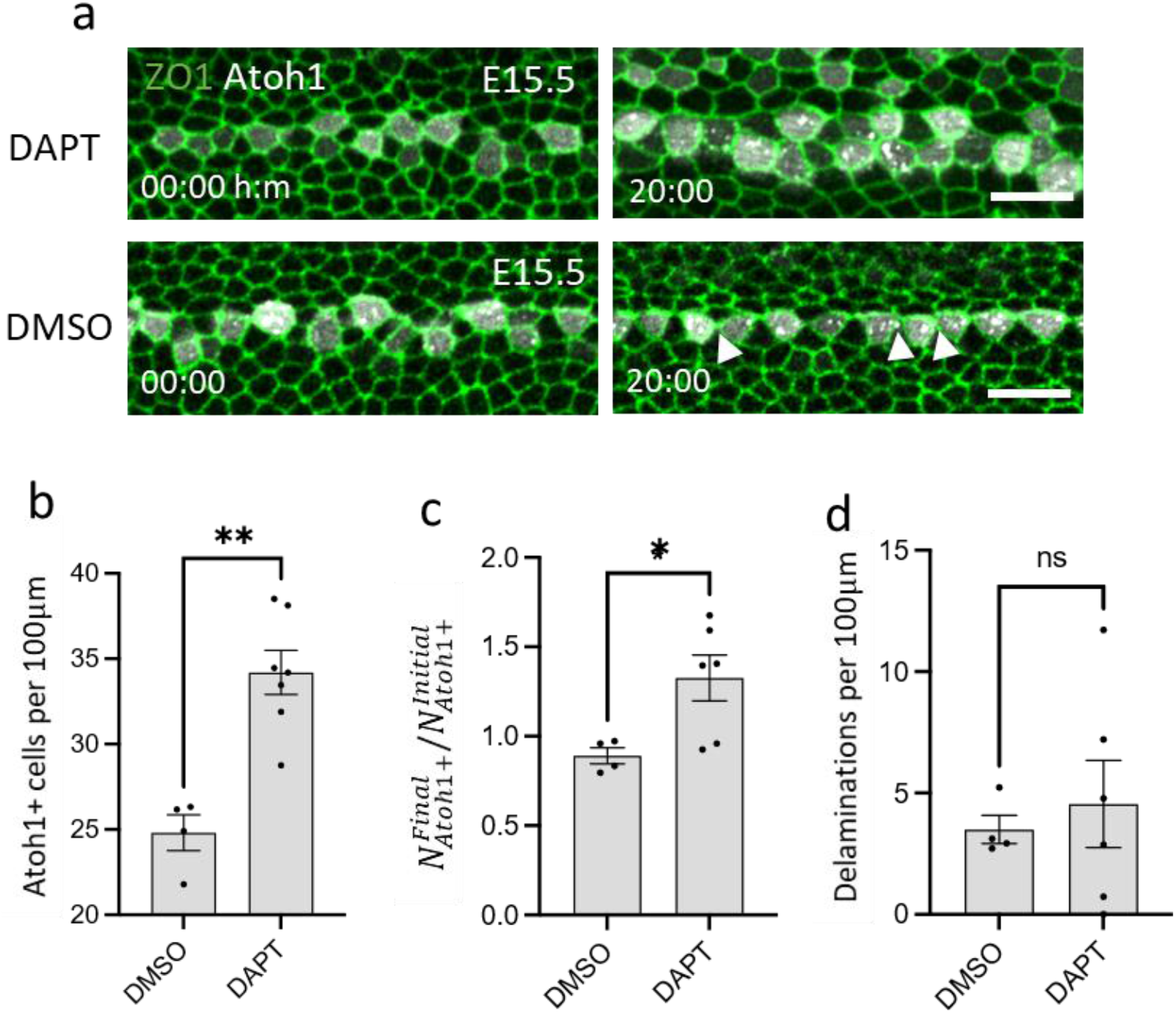
Partial Notch inhibition results in extra IHC. **(a)** Filmstrips from a movie of E15.5 cochlear explants subjected to either 1μM of the Notch inhibitor DAPT (top row, Movie S11) or DMSO control (bottom row). **(b)** Total number of Atoh1^+^ cells per 100μm 24 hours after treatment. **(c)** Ratio between the final (24h after treatment) and initial (0h after treatment) number of Atoh1^+^ cells. **(d)** Number of delaminating Atoh1^+^ cells per 100μm during the movie. Statistics: Mann-Whitney test in (b) and (d), student’s t-test in (c), all plots show mean ± SEM. Repeats in (b-d): n=6, 4 for DAPT and DMSO, respectively. **P<0.01, *P<0.05, ns= not significant. Scale bars in (a): 10μm.

**Figure S5.**
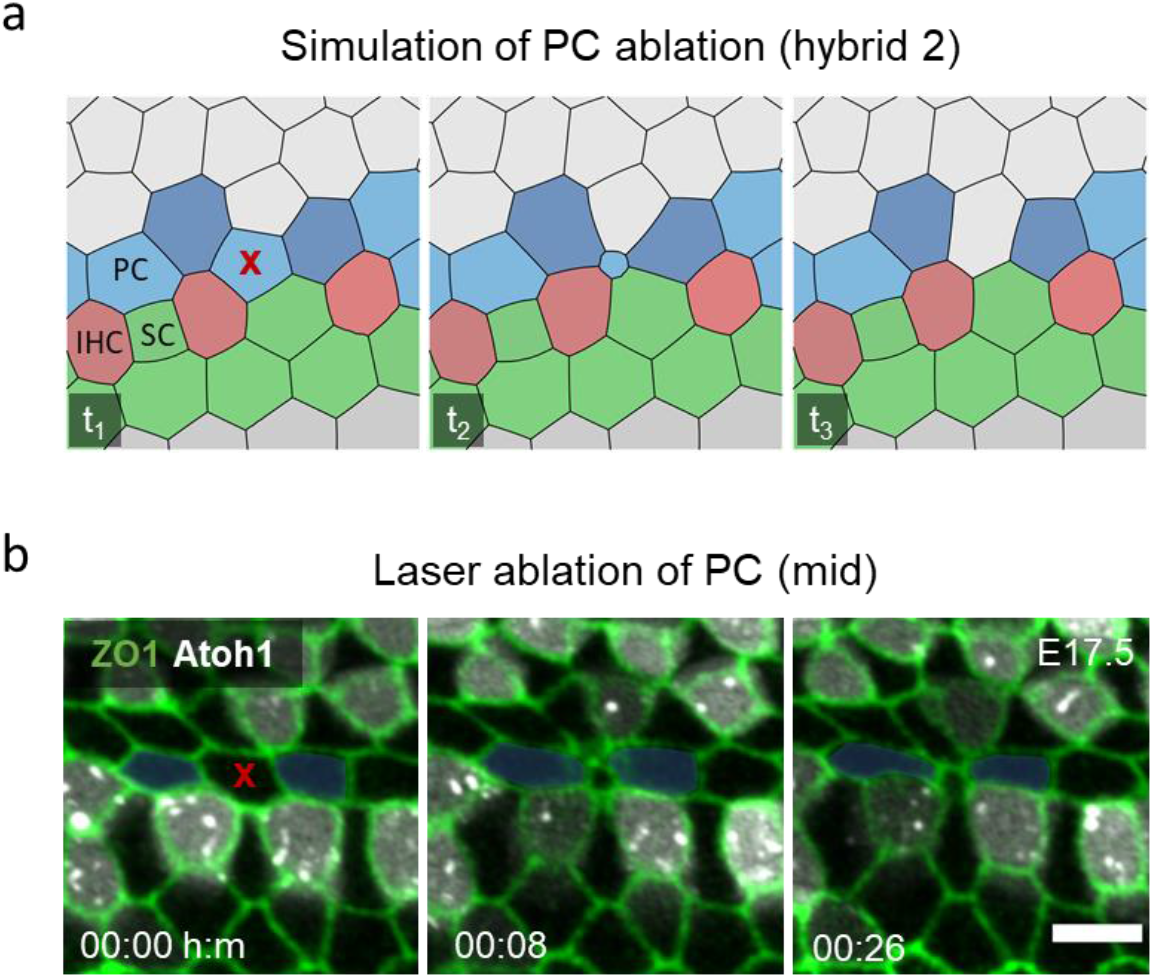
Laser ablation experiments support a preferential adhesion model. **(a)** Filmstrip of a simulation from the hybrid 2 model (without adhesion rules), where a single PC was ablated (red ‘x’) (Movie S14). The gap created by the ablated PC is not repaired by the neighboring PC leaving a gap in the PC row. **(b)** A filmstrip from a laser ablation experiment performed at the mid region of an E17.5 cochlea, where a single PC was ablated (red ‘x’) leaving a gap in the PC row (Movie S16). Scale bar: 5μm

**Figure S6.**
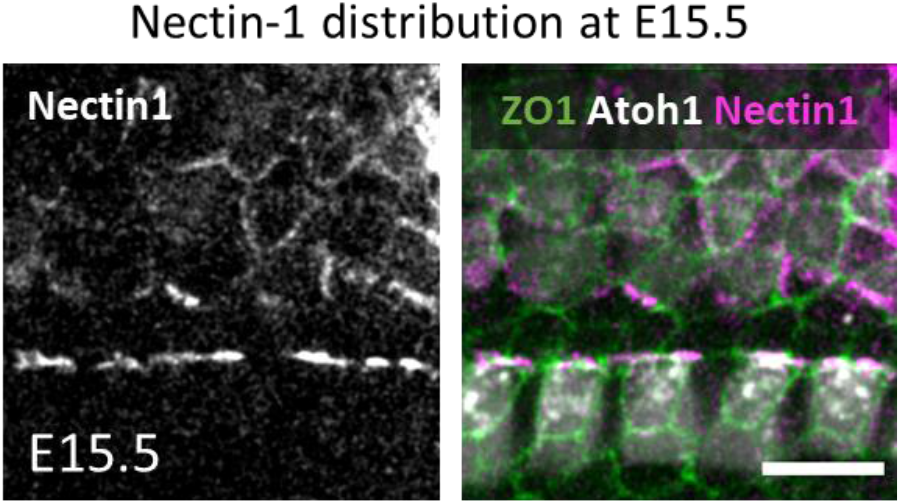
Nectin-1 distribution in the developing OoC. Antibody staining of an E15.5 cochlea with α-Nectin-1 antibody showing localization of Nectin-1 at the IHC:PC boundary. Scale bar: 10μm.

## Movie Captions

**Movie S1. Associated with Fig. 1 in the main text**. A movie from an E15.5 cochlear explant, showing the differentiation wave of HC in the base-to-apex direction. Scale bar: 50μm.

**Movie S2. Associated with Fig. 1d in the main text.** Simulation of the lateral inhibition model. The expression of Notch and its ligand is represented by the green and red hues, respectively.

**Movie S3. Associated with Fig. 1e in the main text**. A movie from the apex region of an E15.5 cochlear explant, showing the differentiation of IHC and their organization into a single line. Scale bar: 10μm.

**Movie S4. Associated with Fig. S1d in the supplementary information.** A movie from a E15.5 cochlea in a medial region to the OoC, showing two cell divisions. Scale bar: 5μm.

**Movie S5. Associated with Fig. S2a in the supplementary information**. A movie from an E15.5 cochleae explant, showing an IHC (red dot) performing intercalation toward the PC row. Scale bar: 5μm.

**Movie S6. Associated with Fig. 2a in the main text**. A movie from an E15.5 cochlear explant, showing an IHC (red dot) performing hopping intercalation toward the PC row. Scale bar: 5μm.

**Movie S7. Associated with Fig. 3b in the main text**. A simulation of an IHC (red) performing hopping intercalation toward the PC row (blue).

**Movie S8. Associated with Fig. 3d in the main text**. A simulation of hybrid 1 model. The expression of Notch and its ligand is represented by the green and red hues, respectively.

**Movie S9. Associated with Fig. 4a in the main text**. A movie from an E15.5 cochleae explant, showing two Atoh1+ cells (red dots) delaminate. Scale bar: 5μm.

**Movie S10. Associated with Fig. 4e in the main text.** A simulation of hybrid 2 model. The expression of Notch and its ligand is represented by the green and red hues, respectively.

**Movie S11. Associated with Fig. S4a in the supplementary information**. A movie from an E15.5 cochlear explant treated with 1μM of Notch inhibitor DAPT (added at 00:00h:m). Scale bar: 10μm.

**Movie S12. Associated with Fig. 5d in the main text**. A simulation hybrid 3 model. This model starts at the final timepoint of hybrid 2 model simulation.

**Movie S13. Associated with Fig. 5g in the main text**. A simulation of a PC ablation, performed at the final timepoint of hybrid 3 model simulation (includes adhesion rules).

**Movie S14. Associated with Fig. S5a in the supplementary information**. A simulation of a PC ablation, performed at the final timepoint of hybrid 2 model simulation (does not include adhesion rules).

**Movie S15. Associated with Fig. 5i in the main text**. A laser ablation experiment performed at the base region of an E17.5 cochlea, where a single PC was ablated. Scale bar: 5μm.

**Movie S16. Associated with Fig. S5b in the supplementary information**. A laser ablation experiment performed at the mid region of an E17.5 cochlea, where a single PC was ablated. Scale bar: 5μm.

